# Murine gut microbiota dysbiosis via enteric infection modulates the foreign body response to a distal biomaterial implant

**DOI:** 10.1101/2025.01.13.632473

**Authors:** Brenda Yang, Natalie Rutkowski, Anna Ruta, Elise Gray-Gaillard, David R. Maestas, Sean H. Kelly, Kavita Krishnan, Xinqun Wu, Shaoguang Wu, Allen Chen, Connor D. Amelung, Joscelyn C. Mejías, Joshua S. T. Hooks, Isabel Vanderzee, Patricia Mensah, Nazmiye Celik, Marie Eric, Peter Abraham, Ada Tam, Sharon Gerecht, Franck Housseau, Drew M. Pardoll, Cynthia L. Sears, Jennifer H. Elisseeff

## Abstract

The gut microbiota influences systemic immunity and the function of distal tissues, including the brain, liver, skin, lung, and muscle. However, the role of the gut microbiota in the foreign body response (FBR) and fibrosis around medical implants is largely unexplored. To investigate this connection, we perturbed the homeostasis of the murine gut microbiota via enterotoxigenic *Bacteroides fragilis* (ETBF) infection and implanted the synthetic polymer polycaprolactone (PCL) into a distal muscle injury. ETBF infection in mice led to increased neutrophil and γδ T cell infiltration into the PCL implant site. ETBF infection alone promoted systemic inflammation and increased levels of neutrophils in the blood, spleen, and bone marrow. At the PCL implant site, we found significant changes in the transcriptome of sorted fibroblasts but did not observe gross ETBF-induced differences in the fibrosis levels after 6 weeks. These results demonstrate the ability of the gut microbiota to mediate long-distance effects such as immune and stromal responses to a distal biomaterial implant.

**Significance Statement:** The foreign body response to implants leads to chronic inflammation and fibrosis that can be highly variable in the general patient population. Here, we demonstrate that gut dysbiosis via enteric infection promoted systemic inflammation and increased immune cell recruitment to an anatomically distant implant site. These results implicate the gut microbiota as a potential source of variability in the clinical biomaterial response and illustrate that the local tissue environment can be influenced by host factors that modulate systemic interactions.

## Main Text Introduction

The development of implantable biomaterials and medical devices has improved quality of life for many patients by enabling the replacement of missing body parts and restoration of tissue function. For instance, breast implants offer reconstructive options for mastectomy patients and artificial hip joints provide improved mobility for those with damaged or injured hips. However, the long-term performance of these implants can be compromised due to complications induced by the foreign body response (FBR), which is characterized by chronic inflammation and fibrosis (1, 2). Fibrotic encapsulation of the implant can cause significant pain, impair implant function, and even necessitate implant removal (3, 4). In the FBR, blood-material interactions cause protein adsorption at the implant surface. This is followed by complement activation and recruitment of innate immune cells, including neutrophils, monocytes, and macrophages. Macrophages fuse into foreign body giant cells that surround the implant and release reactive species in an attempt to phagocytose the implant. Recruited lymphocytes such as T helper 2 (Th2) cells also release factors that promote macrophage fusion (2). Lastly, fibroblasts are activated by the local inflammatory environment to secrete extracellular matrix (ECM) proteins that encapsulate the implant in dense fibrotic tissue. Our understanding of the key cell types and cell-cell interactions that mediate the FBR continues to grow (5). Recent studies have implicated the adaptive immune response, including T helper 17 (Th17) cells and B cells, and stromal cell populations including senescent cells in the FBR to naturally derived and synthetic materials (6, 7). Recognition that the adaptive immune system can contribute to the FBR introduces a potential source of variation in the implant response that is observed clinically.

Biomaterial properties such as composition, size, shape, and stiffness impact host cell recruitment and development of fibrosis. For instance, modification of the physical properties of a biomaterial, addition of a bioactive surface coating, or use of anti-fouling materials that resist protein adsorption can decrease FBR-induced inflammation or improve implant integration (3, 8). However, there is clinical variability in the response to implants and levels of fibrosis across patients, suggesting there are biomaterial-extrinsic factors that modulate the FBR. A number of host factors such as history of infection, diet, age, sex, and the gut microbiota likely contribute to variability in the FBR.

The gut microbiota is emerging as a key player in a variety of health and disease outcomes, including immune modulation, metabolic disorders, cardiovascular diseases, and certain cancers (9). It is comprised of ∼10^14^ microorganisms including bacteria, viruses, fungi, and archaea that colonize the gastrointestinal (GI) tract and is highly variable across the population (10). Gut bacteria co-evolved with humans to form a mutually beneficial relationship in which the intestines provide nutrients to the bacteria while the bacteria aid with digestion of food, strengthen integrity of the gut barrier, and protect against pathogens (10).

The influence of gut bacteria extends beyond the GI tract. For instance, gut bacteria are critical in shaping host immunity (11, 12). Depletion of gut bacteria in mice via broad-spectrum antibiotics or germ-free conditions leads to dysregulation of myeloid cell, lymphocyte, and proinflammatory cytokine levels in GI and systemic sites (11). The gut bacterial composition is also linked to a wide range of immune-related clinical outcomes in mice and humans, including susceptibility to pathogenic enteric infections like *Clostridioides difficile* (13), exacerbation of autoimmune diseases (14), and responsiveness to immune checkpoint inhibitor cancer therapies (15). Not only does the gut microbiota interact with our immune system, but it also communicates with a multitude of peripheral tissues. Studies on a variety of gut-tissue axes such as the gut-brain (16–18), gut-skin (19), gut-muscle (20–22), gut-lung (23, 24), and gut-pancreas (25, 26) axes demonstrate that gut bacteria can influence tissue-specific functions in both homeostatic and pathologic conditions. Though the gut microbiota has a systemic impact on the body and influences the immune response in tissues beyond the GI tract, its role in modulating the immune-driven FBR is largely unknown. One recent study demonstrated that depletion of the gut microbiota reduced host cell infiltration into a subcutaneous silicone implant and decreased thickness of the fibrotic capsule around the implant (27).

Here, we sought to further investigate the gut microbiota-FBR connection using a specific infection model with enterotoxigenic *Bacteroides fragilis* (ETBF). ETBF is a pathogenic strain of the common human commensal *Bacteroides fragilis* that secretes a metalloprotease called *Bacteroides fragilis* toxin (bft) (28). ETBF is associated with diarrheal illnesses, inflammatory bowel disease, and colorectal cancer (28, 29). In mice, ETBF infection induces an acute and chronic Th17 response within the colon (30, 31). We demonstrate that dysbiosis of the gut microbiota via ETBF infection induces a local and systemic inflammatory state, significantly alters muscle-specific gene expression, and heightens the inflammatory response to a distal biomaterial implant. Further, we establish that ETBF infection alters the transcriptome of fibroblasts at the implant site and modulates expression of proteoglycans in the fibrous capsule.

## Results

### ETBF infection promotes inflammation in the GI tract and at distal muscle tissue with biomaterial implant

To model a biomaterial response and the subsequent fibrosis, we performed bilateral volumetric muscle loss (VML) surgery in the quadricep muscles of mice and implanted polycaprolactone (PCL) in the defect space. The injury induced by the VML surgery triggers a potent immune response, and the PCL implant further produces a robust fibrotic and type 3 inflammatory response (6). In order to determine the impact of ETBF infection on the immune response to a PCL implant, we designed an experimental model that coupled these two anatomically distant biological insults (Fig. 1A). We first administered a 4-day course of antibiotics (clindamycin and streptomycin) in the drinking water to establish a favorable environment for ETBF colonization, consistent with previously published protocols (30–32). On the day of inoculation, we returned the mice to the facility’s drinking water and orally gavaged mice with either Dulbecco’s phosphate buffered saline (DPBS) (control) or ∼10^8^ CFU of ETBF suspended in DPBS. We performed VML surgery and PCL implantation 1-week after ETBF inoculation, which corresponds to the acute colitis phase within the mice (32, 33). We conducted most of the tissue readouts at 6- weeks post VML+PCL, as it takes approximately 6 weeks for the fibrotic capsule to form around the PCL implant (6).

**Fig. 1.**
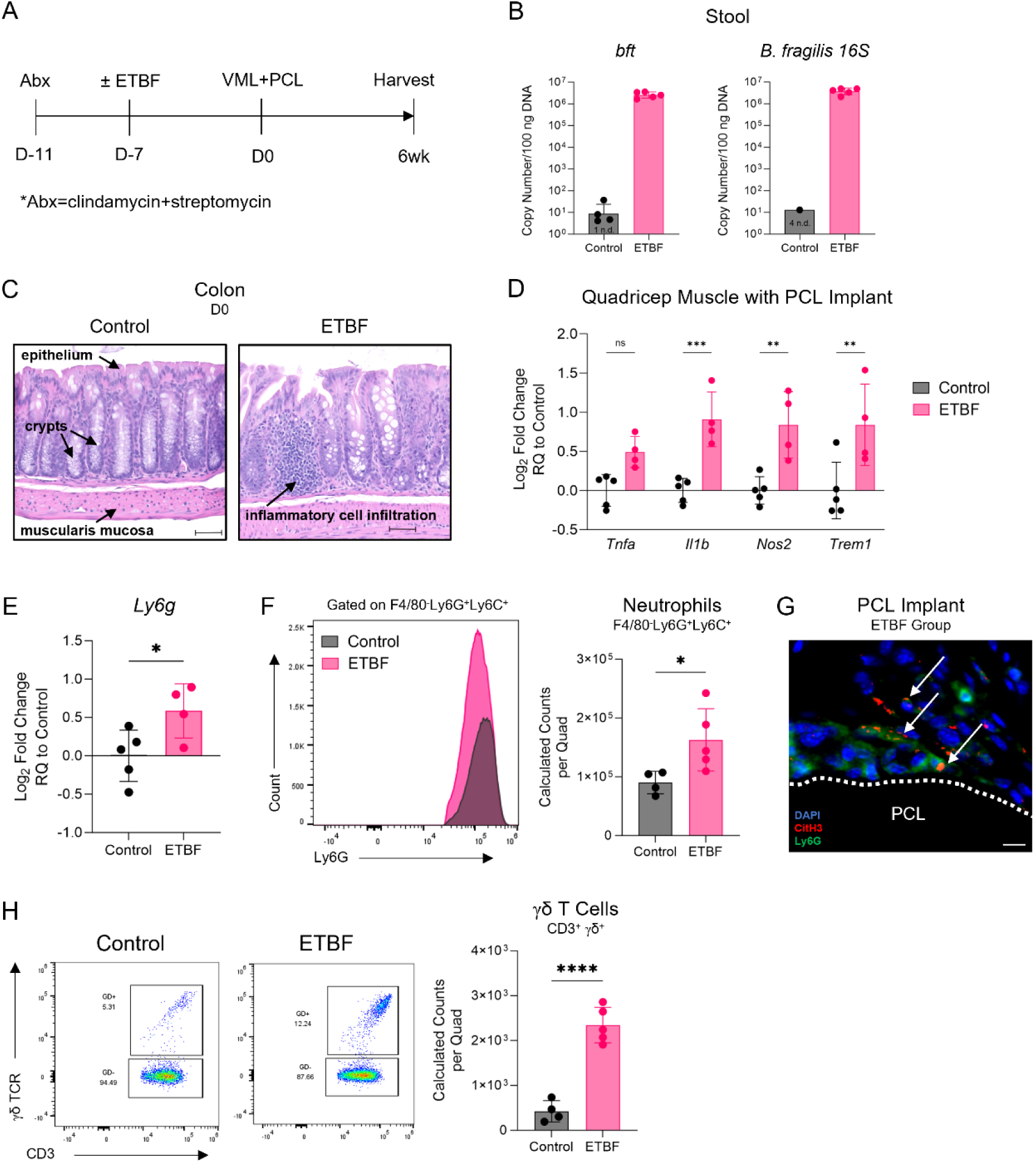
Colonic ETBF infection increases inflammation and immune cell infiltration into PCL implant. (A) Experimental timeline. Abx = antibiotics (clindamycin and streptomycin) (see Methods for details). (B) Stool detection of *B. fragilis* toxin (*bft*) and *B. fragilis 16S* at the 6-week harvest time point. Copy number calculated through standard curve. n.d. = not detected. (C) H&E image of colon at the Day 0 time point (1-week post gavage of DPBS or ETBF). Scale bar, 100 μm. (D) Quadricep muscle with PCL implant expression of inflammatory genes at 6-weeks post VML+PCL in mice gavaged with ETBF, normalized to the control condition. *Rer1* used as reference housekeeper gene. (E) Quadricep muscle with PCL implant expression of *Ly6g* at 6-weeks post VML+PCL in mice gavaged with ETBF, normalized to the control condition. *Rer1* used as reference housekeeper gene. (F) Calculated neutrophil (CD11b^+^F4/80^-^Ly6G^+^Ly6C^+^) counts in quadricep muscle with PCL implant at 6-weeks post VML+PCL in mice gavaged with DPBS or ETBF. (G) Immunofluorescence staining of DAPI (blue, nuclei), citrullinated histone H3 (red, NETs), and Ly6G (green, neutrophils) in PCL implant site of ETBF-infected mouse at 6-weeks post VML+PCL. White arrows point to Ly6G^+^CitH3^+^ cells. Scale bar, 10 μm. (H) Calculated γδ T cell counts in quadricep muscle with PCL implant at 6-weeks post VML+PCL in mice gavaged with DPBS or ETBF. Graphs show geometric mean ± geometric SD (B) or mean ± SD (D-F, H). N=4-5 (B, D-F, H). *P < 0.05, **P < 0.01, ***P < 0.001, and ****P < 0.0001 by two-way ANOVA with Sidak’s multiple comparisons (D) or two-tailed unpaired t test (E-F, H).

First, we confirmed ETBF colonization and the subsequent immune changes in GI tissues. At 1-week post inoculation, high levels of ETBF colonization (10^5^-10^6^ copies/100 ng DNA were observed in the stool of the infected group, with either non-detectable levels or levels below the noise threshold (<10^2^ copies/100 ng DNA) in the uninfected group (SI Appendix, Fig. S1A). Consistent with prior work (30, 33), ETBF induced inflammatory cell infiltration in the colonic mucosa at 1-week post inoculation (Fig. 1C). ETBF infection also increased inflammatory gene expression in the colon, including *Il1b*, *Il6*, and *Ifng* (SI Appendix, Fig. S1B). The mesenteric lymph nodes of ETBF-infected mice displayed similar increases in inflammatory gene expression, in addition to a more prominent type 3 phenotype via significantly increased *Il17a* and *Il17f* (SI Appendix, Fig. S1C). At the 6-week harvest time point, ETBF colonization levels remained high in the infected group and either non-detectable or below-threshold in the control group (Fig. 1B). At the PCL implant site, ETBF infection led to elevated expression of multiple inflammatory genes (Fig. 1D-E), with *Il1b* and *Trem1* suggesting increased myeloid cell activation (34, 35), *Tnfa* and *Nos2* indicative of increased T cell and myeloid cell activity (36, 37), and *Ly6g* relevant to neutrophils.

### ETBF infection increases neutrophil and T cell recruitment to distal PCL implant site

To follow up on the inflammatory changes, we performed spectral flow cytometry on the quadricep muscle with PCL implant at the 6-week time point. ETBF infection significantly increased the numbers of immune cells (CD45^+^) infiltrating the implant site (SI Appendix, Fig. S2A). In concordance with the increased *Ly6g* expression, ETBF infection also significantly increased neutrophil numbers (CD11b^+^F4/80^-^Ly6G^+^Ly6C^+^) (162,884 ± 53,099) compared to non-infected mice (90,237 ± 19,154) (Fig. 1F). To further investigate the kinetics of neutrophil infiltration, we assessed multiple time points and found that implant-associated neutrophils and *Ly6g* expression remained elevated at 3-days and 1-week post VML+PCL, but returned to baseline or below baseline by the 12-week time point (SI Appendix, Fig. S2B, Fig. S2E).

To determine where the neutrophils were localizing and how they were functioning at the implant site, we performed immunofluorescence staining for Ly6G and citrullinated histone H3 (CitH3), a commonly used marker for neutrophil extracellular traps (NETs) (38). NETs are extracellular structures consisting of decondensed DNA, histones, and granule proteins that are released by neutrophils in response to pro-inflammatory stimuli or microbial cues (39). Without proper regulation, NETosis can exacerbate inflammation and contribute to excessive tissue damage (39). At the 6-week time point, we observed that neutrophils localized at the surface of the PCL particles and stained positive for CitH3, confirming activation and secretion of NETs (Fig. 1G).

Due to the increase in *Tnfa* and *Nos2* expression at the implant site (Fig. 1D) as well as the growing appreciation for the role of T cells in the FBR (5), we explored the impact of ETBF infection on the T cell response to the PCL implant. At the 6-week time point, there were significantly more γδ T cells at the implant site of ETBF-infected mice (2344 ± 397) relative to the control uninfected mice (425 ± 236) (Fig. 1H). This significant increase in γδ T cell numbers induced by ETBF was also present at the 3-day and 1-week time points (SI Appendix, Fig. S2C). We found similar trends with CD4 T cells which increased at the implant site in ETBF-infected mice (SI Appendix, Fig. S2D). Taken together, these findings demonstrate that an enteric infection can amplify the innate and adaptive immune response to an anatomically distant biomaterial implant.

### Impact of enteric infection on immune response to distal implant depends on type of biomaterial and type of infection

To determine if the ETBF-induced immune response at the implant was specific to PCL, we investigated the impact of ETBF infection on a variety of biomaterial implants including a biological ECM derived from porcine urinary bladder (SI Appendix, Fig. S3A) and a synthetic polyethylene glycol (PEG)-acrylate (20% wt/v) hydrogel (SI Appendix, Fig. S3A). Using the same experimental timeline as outlined in Fig. 1A, we observed that in contrast to the PCL implant, ETBF infection did not impact the number of infiltrating neutrophils or γδ T cells at the ECM or PEG implant site (SI Appendix, Fig. S3B). In addition, ETBF infection led to lower numbers of overall immune cells (CD45^+^) and significantly lower numbers of CD4 T cells at the ECM implant site (SI Appendix, Fig. S3B), opposite to what was observed at the PCL implant site (SI Appendix, Fig. S2A, Fig. S2D). These results demonstrate that ETBF can induce notably different immune responses at the implant site depending on the type of biomaterial implant.

In addition to the ECM and PEG implants, we tested the impact of ETBF infection on the immune response to two clinical HA hydrogels (SI Appendix, Fig. S4A). We altered the experimental timeline to more closely recapitulate the clinical scenario in which patients receive the HA hydrogel implant and subsequently acquire an enteric infection (SI Appendix, Fig. S4A). Though ETBF infection led to a trending increase in *Tnfa* expression in the PCL implant site (Fig. 1D), it did not induce changes in *Tnfa* levels at the HA hydrogel implant sites (SI Appendix, Fig. S4B). To investigate beyond the local implant site, we analyzed regional changes at the draining inguinal lymph nodes, as lymph nodes are sites of immune surveillance that promote immune cell activation and mobilization during inflammation (40). Interestingly, even with an altered experimental timeline, ETBF infection significantly increased γδ T cell numbers in the inguinal lymph nodes of mice implanted with HA hydrogel 2 (SI Appendix, Fig. S4C). The ETBF-induced increase in γδ T cells in the PCL implant site and in the lymph nodes draining the HA hydrogel implant site suggests that γδ T cells may be important regulators of communication between the dysbiotic gut and a distal biomaterial implant.

To evaluate whether the type of enteric infection altered the immune response at the implant site, we utilized an infection model with a strain of *Escherichia coli* (*E. coli*) that carries the pathogenicity island polyketide synthase (*pks*). *Pks* encodes the genotoxin colibactin, which induces DNA damage in colonic epithelial cells (41). We followed a similar experimental timeline as described in Fig. 1A, except cefoxitin was used to facilitate colonization with this strain of *E. coli* (Methods; SI Appendix, Fig. S5A) (41). Following 1-day of antibiotic (cefoxitin) washout, we gavaged control mice with DPBS while treatment mice received *E. coli* suspended in DPBS. At the 6-week harvest time point, we observed moderate levels of *E. coli* colonization in the stool of the infected group (∼10^4^ copies/100 ng DNA) and non-detectable levels in the control group (SI Appendix, Fig. S5B). Notably, mice with *E. coli* had 2-3 orders of magnitude lower levels of colonization than mice with ETBF at the 6-week harvest time point (∼10^4^ copies/100 ng DNA vs. 10^6^-10^7^ copies/100 ng DNA) (SI Appendix, Fig. S5B, Fig. 1B). Upon profiling the immune landscape at the PCL implant site, we observed that *E. coli* infection significantly increased the frequency of γδ T cells (SI Appendix, Fig. S5C), similar to what had been seen with ETBF infection (SI Appendix, Fig. S2A). However, *E. coli* infection induced a significant decrease in CD4 T cell frequency and a slight trending increase in neutrophil frequency (SI Appendix, Fig. S5C). Therefore, the immune response at the distal PCL implant site varied depending on the enteric infection, providing further evidence of the gut microbiota serving as a source of variability in the immune response to implants.

### ETBF-infected mice display systemic inflammation and amplified neutrophil trafficking

Because ETBF infection had consistently high levels of colonization and induced robust inflammatory changes at the PCL implant site, we further investigated this model. We sought to determine how ETBF infection altered the systemic immune homeostasis of the mice prior to the secondary insult of the distal PCL implant. We treated the mice with antibiotics (clindamycin and streptomycin) for 4- to 5-days, inoculated with DPBS or ETBF, and evaluated changes at the 1- week time point post inoculation (Fig. 2A). ETBF infection increased inflammatory gene expression in the spleen, including *Il1b* and *Chil3* (Fig. 2B). *Chil3* is commonly used as an M2-macrophage marker, but is also expressed by neutrophils (42) and is associated with neutrophilia in nematode infection models (43, 44). In combination with the elevated *Ly6g* expression (Fig. 2B), these results suggest that ETBF infection may increase migration of neutrophils to secondary lymphoid tissues such as the spleen.

**Fig. 2.**
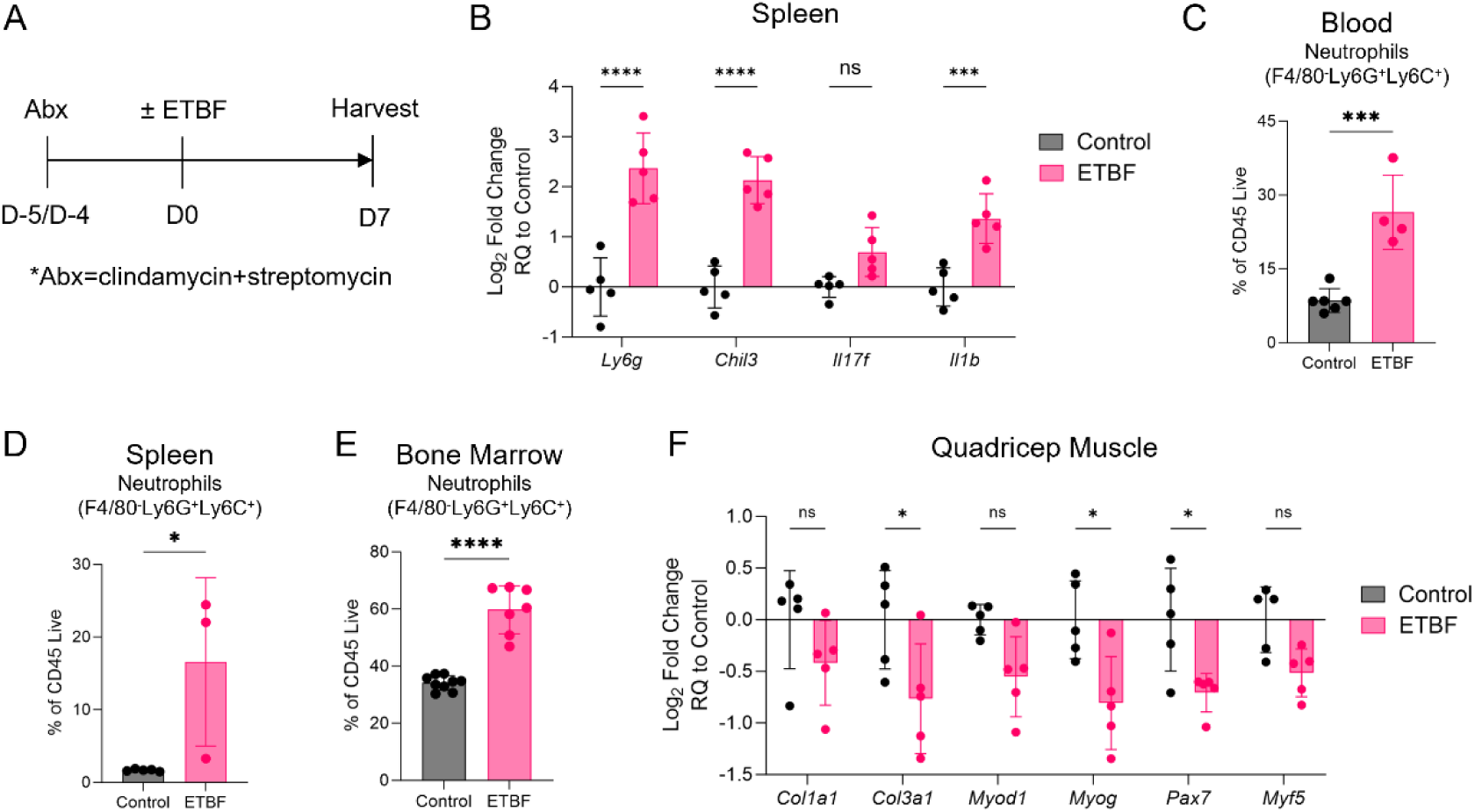
ETBF infection impacts systemic immune and non-immune tissues. (A) Experimental timeline. (B) Spleen expression of inflammatory genes at 1-week post ETBF, normalized to the control condition. *B2m* used as reference housekeeper gene. (C) Neutrophil (CD11b^+^F4/80^-^Ly6G^+^Ly6C^+^) percentage of CD45^+^ cells in blood at 1-week post gavage. (D) Neutrophil (CD11b^+^F4/80^-^Ly6G^+^Ly6C^+^) percentage of CD45^+^ cells in spleen at 1-week post gavage. (E) Neutrophil (CD11b^+^F4/80^-^Ly6G^+^Ly6C^+^) percentage of CD45^+^ cells in femur bone marrow at 1-week post gavage. (F) Quadricep muscle expression of muscle and ECM-related genes at 1- week post ETBF, normalized to the control condition. *Rer1* used as reference housekeeping gene. Graphs show mean ± SD (B-F). N=3-5 for all graphs (B-F). *P < 0.05, **P < 0.01, ***P < 0.001, and ****P < 0.0001 by two-way ANOVA with Sidak’s multiple comparisons (B, F) or two-tailed unpaired t test (C-E).

To further investigate the immune changes, we used flow cytometry to evaluate the levels of neutrophils in the circulation and lymphoid tissues. We observed a significantly higher frequency of neutrophils in the blood of ETBF-infected mice (26.5 ± 7.56% of CD45^+^) compared to control mice (8.58 ± 2.42% of CD45^+^) (Fig. 2C). Additionally, ETBF-infection significantly increased neutrophil frequencies in the spleen (16.59 ± 11.58% of CD45^+^ vs. 1.68 ± 0.15% of CD45^+^) and bone marrow (59.7 ± 8.34% of CD45^+^ vs. 34.2 ± 2.45% of CD45^+^) (Fig. 2D-E). These results support the notion that ETBF infection induces granulopoiesis in the bone marrow and promotes mobilization of neutrophils to secondary sites via the circulation. When treating mice with both ETBF infection and a PCL implant, we observed that at the earlier time points, ETBF led to significant increases in neutrophil frequencies in the blood (3-day and 1-week time point), spleen (1-week time point), and bone marrow (3-day time point) (SI Appendix, Fig. S6A-C). At the 6-week time point, neutrophil frequencies remained elevated in the bone marrow of ETBF-infected mice (SI Appendix, Fig. S6C). Given that the ETBF-infected mice exhibited increased levels of neutrophils at the implant site itself (Fig. 1F, SI Appendix, Fig. S2B), these findings indicate that neutrophils contribute to immune communication between the dysbiotic gut microbiota and a distal biomaterial implant.

### ETBF infection reduces expression of collagens and myogenic markers in naïve quadricep muscle

Since the ETBF infection induced systemic immune changes, we evaluated how ETBF infection affected the gene expression within the naïve quadricep muscle (with no injury or implant). At 1-week post inoculation, ETBF infection led to reduction in the quadricep muscle expression of fibrillar collagens (*Col1a1*, *Col3a1*) (Fig. 2F), which are ECM components that confer tensile strength and elasticity to muscle tissue (45). ETBF infection also reduced expression of genes associated with satellite cell activation (*Pax7*) and myogenic differentiation (*Myod1, Myog, Myf5*) in the quadricep muscle (Fig. 2F), which are crucial for skeletal muscle regeneration after injury (46). These results demonstrate that a local enteric infection can influence homeostasis of distal non-immune tissues such as the quadricep muscle.

### ETBF infection impacts ECM composition-related transcriptome of fibroblasts within the PCL implant

After observing that ETBF infection increased immune cell infiltration into the implant site, we sought to determine if this corresponded to changes in fibrosis levels around the implant. The relative red-green ratio in Picrosirius Red images is often used as a metric of fibrosis, with higher ratios indicating more of the “thick” collagen fibers relative to “thin” fibers (47). Quantification of Picrosirius Red-stained sections of the implant site did not result in any differences between the two groups at the 6-week time point (Fig. 3A, SI Appendix, Fig. S7). There were no notable qualitative differences in the fibrosis around the PCL particles via Masson’s Trichrome staining between the control and infected groups at the 6-week (SI Appendix, Fig. S8A) or 12-week (SI Appendix, Fig. S9A) time point. In addition, ETBF infection did not impact expression of fibrotic genes including *Col1a1*, *Col3a1*, and *Tgfb1* within the PCL implant site at the 6-week (SI Appendix, Fig. S8B) or 12-week (SI Appendix, Fig. S9B) time point.

**Fig. 3.**
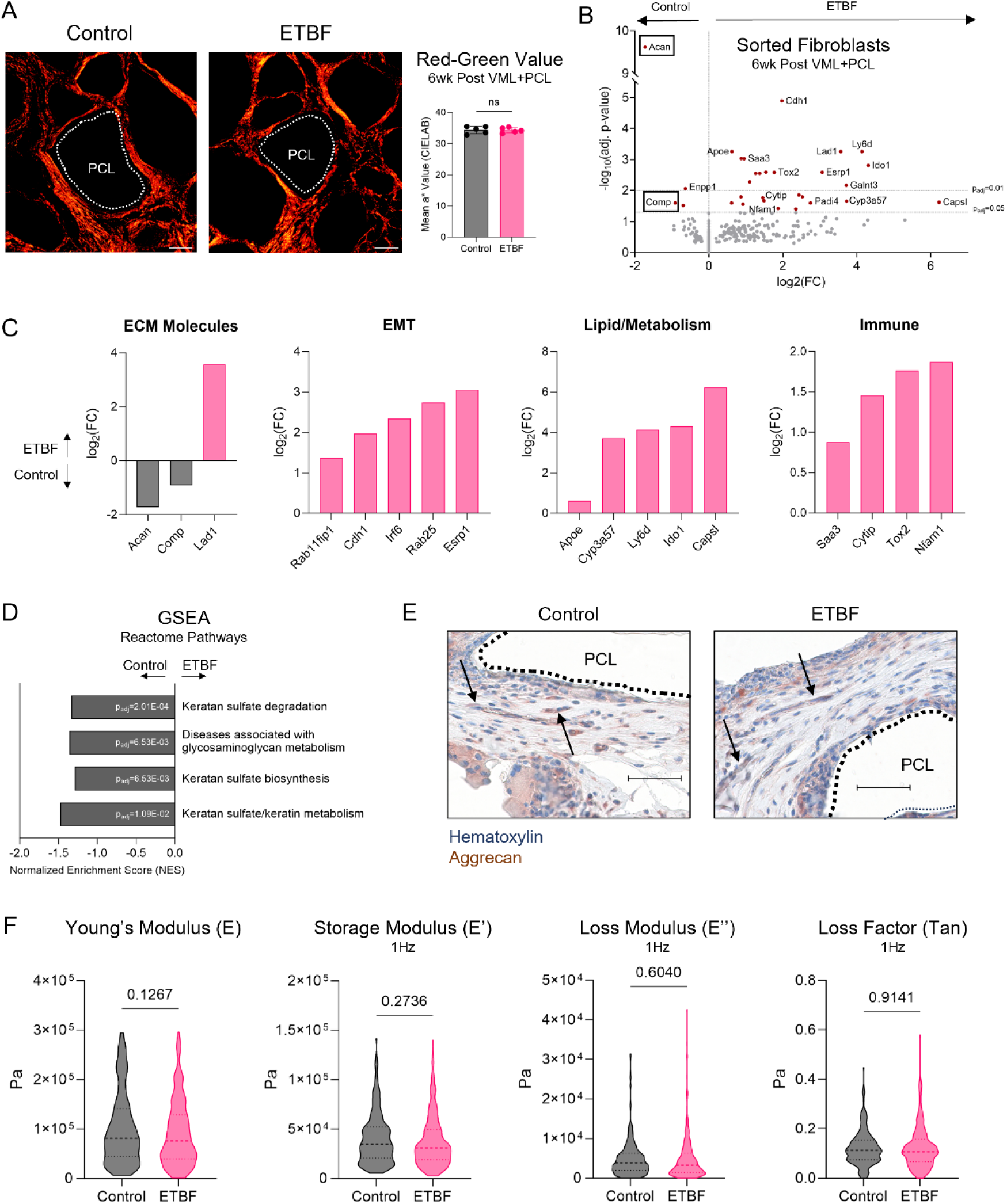
ETBF infection alters the transcriptional profile of fibroblasts at the PCL implant site. (A) Picrosirius Red images of PCL implant site at 6-weeks post VML+PCL in mice gavaged with DPBS or ETBF. Scale bar, 100 μm. Quantification of mean red-green value on CIELAB axis shown on right. (B) Volcano plot of differential gene expression results of bulk RNA sequencing of sorted fibroblasts (CD45^-^CD31^-^CD29^+^) from PCL implant site at 6-weeks post VML+PCL. (C) Based on volcano plot in (B), manual categorization of significant (p_adj_ < 0.05) differentially expressed genes of sorted fibroblasts (CD45^-^CD31^-^CD29^+^) from PCL implant site at 6-weeks post VML+PCL. (D) Normalized enrichment scores and adjusted p values of selected Reactome pathways from ranked gene set enrichment analysis. (E) Immunohistochemistry images of aggrecan in PCL implant site at 6-weeks post VML+PCL in mice gavaged with DPBS or ETBF. (F) Young’s modulus (E), storage modulus (E’) at 1Hz, loss modulus (E’’) at 1Hz, and loss factor (Tan) at 1Hz of fibrotic tissue around PCL particles in control and ETBF-infected mice at 6-weeks post VML+PCL. Graphs show mean ± SD (A) or median and quartiles (F). N=5 (A). Biological n=5, technical n=192-320 (F). (See Methods.) *P < 0.05, **P < 0.01, ***P < 0.001, and ****P < 0.0001 by two-tailed unpaired t test (A, F). NES normalized to size of gene set (D).

We then investigated the transcriptome of the fibroblasts from the implant site by sorting CD45^-^CD31^-^CD29^+^ cells at the 6-week time point when there is significant fibrosis. We performed bulk RNA sequencing on the sorted cells and found 30 significant differentially expressed genes (DEGs) between the infected and non-infected group (Fig. 3B). The DEGs represented genes associated with ECM, epithelial-mesenchymal transition (EMT), lipid/metabolism, and immune response (Fig. 3C, SI Appendix, Fig. S10). We observed a 75-fold increase in *Capsl* in the fibroblasts of ETBF-infected mice relative to control mice (Fig. 3C). *Capsl* has been associated with adipogenesis in human adipocytes (48) and angiogenesis in human retinal endothelial cells (49), but its function in non-endothelial stromal cells is less clear and may be worthy of further investigation. Interestingly, we found that several of the DEGs were commonly associated with cartilage tissue, such as *Acan* (which encodes aggrecan) and *Comp* (which encodes cartilage oligomeric matrix protein). Furthermore, ranked gene set enrichment analysis of the Reactome pathways revealed that multiple of the significant pathways were relevant to glycosaminoglycans (GAGs) including keratan sulfate (Fig. 3D), which are found on aggrecan molecules and lend cartilage its compressive strength.

### ETBF infection influences stiffness of fibrotic tissue around distal implant

Because *Acan* was the most significant DEG, we performed immunohistochemistry staining to confirm protein-level production of aggrecan within the implant site. Aggrecan-positive staining occurred in elongated, fibroblast-like cells in the ECM-heavy region between PCL particles (Fig. 3E). Due to the important role of aggrecan in the mechanical properties of cartilage tissue, we sought to determine whether the ETBF-induced downregulation of *Acan* in the implant-associated fibroblasts altered the mechanical properties of the fibrotic tissue. We performed atomic force microscopy (AFM) on the fibrotic tissue around the PCL particles and observed that ETBF-infected mice had a trending decrease (p=0.1267) in the Young’s modulus (E) relative to control mice (Fig. 3F). This suggests a possible link between fibrotic tissue stiffness and fibroblast *Acan* expression. There were no significant differences in the storage modulus (E’), loss modulus (E’’), or loss factor (Tan) at a 1Hz frequency between the control and ETBF-infected mice (Fig. 3F).

## Discussion

The gut microbiota influences a variety of distal biological processes, including skeletal muscle tissue regeneration (20, 50), nociceptive and stress responses (51), and the pathophysiology of many conditions such as diabetes, obesity, non-alcoholic fatty liver disease, and cardiovascular disease (9). Recent evidence demonstrates that its influence can extend to the FBR (27). The results in our study support the immunological connection between the host gut microbiota and the biomaterial response. We report that an enteric infection with ETBF exacerbates the inflammatory response to a distal PCL implant. At the implant site, ETBF infection led to increased neutrophil counts and *Ly6g* expression until the 6-week time point (Fig. 1E-F, SI Appendix, Fig. S2B, Fig. S2E). In homeostatic conditions, neutrophils are typically cleared from sites of inflammation within a few days via apoptosis and subsequent phagocytosis by macrophages (52, 53). In contrast, chronic neutrophil presence is often a sign of an unresolved response to tissue damage or chronic inflammation (54). Therefore, the increase in implant-associated neutrophils at the 6-week time point suggests that ETBF infection amplifies the chronic inflammatory environment at the PCL implant and delays resolution. ETBF infection also increased implant-associated γδ T cells until the 6-week time point (Fig. 1H, SI Appendix, Fig. S2C). γδ T cells can play both protective and pathogenic roles in disease and infection (55). Their elevated numbers may indicate increased activation and production of inflammatory mediators or may suggest recruitment to tamp down the inflammatory response at the implant site. Future studies in mice that are depleted of neutrophils or γδ T cells could elucidate the individual role of these cell types in the FBR.

The magnitude and composition of the FBR depends on a variety of factors, including the tissue into which the material is implanted (27, 56) and the material properties of the implant itself (3, 27). In our own dual-insult experimental models, we observed that the type of biomaterial implant and type of enteric infection influenced the infection-induced immune changes at the implant site (SI Appendix, Fig. S3-S5). Interestingly, the infection-induced increase in γδ T cells at/near the implant site remained persistent across multiple biomaterials (PCL, HA hydrogel 2) (Fig. 1H, SI Appendix, Fig. S4C) and enteric infections (ETBF, *E. coli*) (Fig. 1H, SI Appendix, Fig. S5C). These results implicate γδ T cells as sensitive responders to enteric infection, biomaterial implantation, or both, thus highlighting an interesting area for further exploration.

In addition, the timing of the enteric infection relative to biomaterial implantation may influence the immune responses seen at/near the implant. For the majority of the experiments, we colonized mice with enteric bacteria prior to implanting a biomaterial (Fig. 1A), which more closely mimics the scenario in which patients with prior enteric infections receive medical implants. In contrast, we also implanted biomaterials and allowed the FBR to develop for several weeks before infecting mice with bacteria (SI Appendix, Fig. S4A). This represents the case in which patients who had received implants were subsequently exposed to enteric bacteria and acquired an infection. It is unclear whether the type of biomaterial or the difference in experimental timeline exerted a greater effect on the ETBF-induced immune response to the HA hydrogel implant. The impact of the timing of each individual biological insult could be a worthwhile topic for future investigation.

The gut microbiota also influences systemic host immunity. Depletion of murine gut bacteria via antibiotic treatment can increase the rate of neutrophil apoptosis in the bloodstream (57–59), diminish the migratory capacity of monocytes (60), reduce common lymphoid progenitors in the bone marrow (61, 62), and decrease total lymphocytes in the blood (61). Disruption of native murine gut microbiota via enteric infection can reduce systemic immune function and promote pathological immune outcomes. Specifically, infection with *Clostridioides difficile* induced loss of medullary and cortical junctions in the thymus, loss of germinal centers in the lymph node, and damage to the kidneys (63), exposure to *Helicobacter hepaticus* promoted mammary carcinoma (64), and infection with *Citrobacter rodentium* led to islet-specific autoimmunity (65). Treatment with probiotic strains, on the other hand, can induce positive systemic effects, such as anti-inflammatory effects in the plasma and lungs (66). Here, we show that ETBF infection promotes systemic inflammation via increased inflammatory gene expression in the spleen (Fig. 2B) and increased neutrophil frequencies in the blood, spleen, and bone marrow (Fig. 2C-E). ETBF infection also decreased satellite cell and myogenic marker gene expression within the quadricep muscle (Fig. 2F), which suggests a potential reduction in the regenerative properties of the muscle. This dysregulation of homeostasis in both immune and non-immune tissues likely predisposes the host towards an exacerbated response to an additional biological insult such as the PCL implant, thus demonstrating the interconnected nature of anatomically distant biological insults.

Previous studies demonstrated a link between the gut microbiota and fibrotic tissue thickness around biomaterial implants (27) or distal organ fibrosis (67). However, we observed that ETBF infection did not influence fibrosis levels at the PCL implant site (Fig. 3A, SI Appendix, Fig. S8-S9). Interestingly, ETBF infection altered the transcriptome of implant-associated fibroblasts (Fig. 3B-D) and slightly decreased the stiffness of the fibrotic tissue (Fig. 3F), demonstrating a potentially novel connection between enteric infection, the stromal response to a biomaterial implant, and the mechanical properties of the fibrotic tissue around the implant. Further studies with different enteric infections, more comprehensive mechanical testing on fresh tissues, and additional independent experimental replicates would strengthen this connection.

There are a variety of methods to assess the impact of the gut microbiota on a distal response, including broad-spectrum antibiotics to reduce bacterial populations (11), germ-free mice that are devoid of any microbes (11), fecal microbiota transplantation in which a complex microbiota is inoculated (68), monocolonization in which a single microbe is inoculated into germ-free mice (69), or infection models in which bacterial species are inoculated into mice that possess native gut microbiota (70). Germ-free mice have dysfunctional immune systems, require high costs and resources for maintenance, and lack microbes at all mucosal and barrier tissues which makes it difficult to causally link the gut microbiota to an outcome of interest (11). On the other hand, broad-spectrum antibiotic treatment allows for studies in mice with intact immune systems and (somewhat) isolates the gut microbiota as a causative factor. However, one of the most commonly used antibiotic treatments (a cocktail of ampicillin, vancomycin, neomycin, and metronidazole) can promote outgrowth of fungal species (71) and induce dehydration and weight loss due to low palatability (72). Monocolonization models can directly evaluate the effect of a specific bacterial species but require that the species is culturable and can survive within the new host. In our study, we utilized an infection model with ETBF, as it does not require the use of specialized gnotobiotic mice, enables the usage of a bacterial species with simple culturing and maintenance, and produces a known immune response within the GI tract. Future studies utilizing additional models of gut dysbiosis would further support the link between the gut microbiota and the FBR.

All of the experiments in this study were performed in young (5- to 6-week old) female mice to eliminate other confounding host factors such as age or sex. Moreover, we conducted these studies in mouse models, which differ significantly from human biology in aspects such as metabolic rate, gut microbiota composition, susceptibility to pathogens, and cognitive development (5). Clinical data can uncover connections between implant-associated adverse events and gut microbiota composition. Lastly, these studies investigated the connection between the gut microbiota and FBR-induced fibrosis, but fibrosis can occur in a variety of pathological contexts including idiopathic pulmonary fibrosis (73), osteoarthritis (74), myocardial disease (75), and age (76). Therefore, a deeper understanding of how the gut microbiota influences fibrotic processes may enable design of new therapeutics targeting tissue fibrosis more broadly.

In conclusion, this work shows that gut dysbiosis via an enteric bacterial infection can have long-lasting consequences on the FBR to a distal biomaterial implant. These findings suggest that the composition of the gut microbiota and previous history of enteric infection should be considered when patients undergo surgeries for medical implants or when implant-associated adverse events occur. Synergizing the effects of host factors and material properties of the implant could potentially reduce the FBR in a much broader patient population.

## Materials and Methods

### Mice

Mice were housed and maintained in the Johns Hopkins Cancer Research Building animal facility in compliance with ethical guidelines outlined by the Johns Hopkins Animal Care and Use Committee (ACUC). All animal procedures were performed in accordance with an approved Johns Hopkins ACUC protocol. 5-6 week old female C57BL/6J mice from Jackson Laboratory (Strain #000664) were used for all experiments.

### Bacteriology and mouse colonization

#### ETBF

All experiments utilized the ETBF strain 86-5443-2-2 (secretes BFT-2) (33), which was kindly provided by the laboratory of Dr. Cynthia Sears at Johns Hopkins School of Medicine. All culture steps listed below were performed in an anaerobic chamber (Model #857-OTA, Plas Labs) with 90% N_2_, 5% H_2_, and 5% CO_2_ at 37°C. ETBF was grown for 2 days on brain-heart infusion (BHI)/agar plates with 37 g/L BHI (BD Bacto), 1.5% agar (BD Bacto), 5 g/L yeast extract (BD Bacto), 50 mg/L L-cysteine (Sigma), 0.5 mg/L hemin (Sigma), 0.1 mg/L vitamin K1 (Sigma), and 10 μg/L clindamycin (Sigma). Isolated colonies were cultured for 2 days in 5 mL BHI broth (same components and concentrations as above, excluding the agar). Liquid culture was subcultured at a 1:100 ratio in 10 mL BHI broth overnight. Bacteria were spun down at 13,000 rpm for 2 minutes and washed twice with 1X Dulbecco’s phosphate buffered saline (DPBS) (Gibco) at 13,000 rpm for 1 minute. Prior to inoculation, mice were treated with 100 mg clindamycin (Sagent Pharmaceuticals or GoldBio) and 5 g/L streptomycin (Sigma) in the drinking water for 4-5 or 7 days. ETBF strain 86-5443-2-2 is resistant to both clindamycin and streptomycin. On the day of inoculation, antibiotics were removed and mice were returned to the facility’s drinking water. Mice were gavaged with 100 μL sterile 1X DPBS (control group) or 100 μL ETBF (∼10^9^ CFU/mL) resuspended in 1X DPBS. The CFUs in the ETBF gavage solution were counted after plating serial dilutions on BHI/agar plates (same as above) and culturing anaerobically at 37°C for 2 days. CFU counts for the ETBF gavage solution were calculated for some but not all experiments.

#### E. coli

*Pks^+^ E. coli* strain NC101 (expressing a fluorescent ampicillin resistance plasmid sfGFP-pBAD) (41) was kindly provided by the laboratory of Dr. Cynthia Sears at Johns Hopkins School of Medicine (originally obtained from Dr. Christian Jobin, University of Florida College of Medicine), with the assistance of Xinqun Wu and Shaoguang Wu. *E. coli* was cultured overnight or for several hours in LB broth with ampicillin. Bacteria were spun down at 13,000 rpm for 2 minutes and washed twice with 1X Dulbecco’s phosphate buffered saline (DPBS) (Gibco) at 13,000 rpm for 1 minute. Prior to inoculation, mice were gavaged with 200 μL of 0.5 g/L cefoxitin (MWI Veterinary Supply) and treated with 0.5 g/L cefoxitin in the drinking water for 2 days. *E. coli* strain NC101 is resistant to cefoxitin. Cefoxitin bottles were removed and mice were returned to the facility’s drinking water for 1 day prior to *E. coli* inoculation. On the day of inoculation, mice were gavaged with 200 μL sterile 1X DPBS (control group) or 200 μL E. coli resuspended in 1X DPBS.

### Volumetric muscle loss (VML) surgery and biomaterial implantation

Bilateral defects in the quadricep muscles of mice were created as previously described (6). Briefly, mice were placed under 2.5% isoflurane for induction of anesthesia and 2% isoflurane for maintenance. 5 mg/kg of an NSAID (Rimadyl, MWI Animal Health) was injected subcutaneously for pain management. The skin above the quadricep muscle was shaved and sprayed with 70% ethanol. A ∼1 cm long incision was made in the skin and the fascia above the quadricep muscle. A ∼3 mm x 4 mm x 4 mm segment of quadricep muscle was resected, and the remaining defect space was filled with a 1.5 mm depth surgical spoon (Roboz) containing one of the following: (1) one leveled scoop of UV-sterilized PCL particles (Mn = 50,000 g/mol, mean particle size <600 μm, Polysciences), (2) 50 μL of 200 mg/mL MicroMatrix UBM Particulate (Integra) in 1X Dulbecco’s phosphate buffered saline (DPBS) (Gibco), (3) 25 μL of 20% wt/v PEG-acrylate hydrogel (see below for details regarding preparation), or (4) 50 μL of Juvèderm Volbella (HA Hydrogel 1) (Allergan) or 50 μL of Juvèderm Ultra 3 (HA Hydrogel 2) (Allergan). Following implantation, the skin was closed with sterile 7 mm wound clips (Roboz).

20% wt/v PEG-acrylate hydrogel was prepared for implantation the day before surgery. A 0.4 g/mL solution of 4-arm PEG-acrylate (MW 20kDa) (Laysan Bio) was prepared in 1X DPBS (Gibco). The solution was heated to 60°C for 20-30 minutes to help dissolve the PEG-acrylate powder. A 50 mg/mL solution of lithium phenyl-2,4,6-trimethylbenzoylphosphinate (LAP) (Tokyo Chemical Industry) was prepared in 1X DPBS (Gibco) and heated to 60°C for 5-10 minutes to help dissolve the LAP. The PEG-acrylate and LAP solutions were mixed at a 1:1 ratio, transferred to a biosafety cabinet, and filtered through a 0.2 μm filter. 25 μL of the mixture was pipetted into molds (caps of 0.2 mL PCR tubes) and photocrosslinked under 405 nm LED light for 45 seconds. PEG-acrylate hydrogels were stored in molds in a sterile container at 4°C overnight prior to implantation.

### Stool collection, DNA isolation, qPCR, and quantification

Stool was collected from the colons of mice, frozen on dry ice, and stored at -80°C. DNA was isolated using either the Quick DNA Fecal/Soil Microbe Miniprep Kit (Zymo) or the QIAamp Fast DNA Stool Mini Kit (Qiagen). For both kits, ∼6 ceramic beads (2.8 mm; OMNI International) were added to the sample tube in the first step and samples were homogenized with a Bead Ruptor 12 (OMNI International) at the highest speed for 15 seconds. The rest of the DNA isolation steps were performed according to manufacturer protocol. DNA concentration was quantified using a NanoDrop 2000 (ThermoFisher Scientific) that was blanked with the appropriate elution buffer depending on the DNA isolation kit used. For the qPCR reaction, 100 ng of stool DNA was added to each well, and TaqMan Gene Expression Master Mix (Applied Biosystems) was added according to manufacturer instructions. A standard curve was generated for each qPCR run using serial dilutions of DNA isolated from liquid ETBF or *E. coli* culture (QIAamp DNA Mini Kit, Qiagen). Expression of *bft* (*Bacteroides fragilis* toxin), *B. fragilis 16S*, *pks*, and *E. coli 16S* was quantified according to the standard curve. SI Appendix, Table S1 lists the primer and probe sequences for *bft*, *B. fragilis 16S*, *pks*, and *E. coli 16S*.

### Tissue collection, processing, qRT-PCR, and analysis

Harvested tissues were immediately placed into either RNAlater (Sigma) or TRIzol (Invitrogen). If placed into RNALater, tissues were stored at 4°C for at least 24 hours before transferring to TRIzol and storing at -80°C. If placed directly into TRIzol, tissues were frozen on dry ice and stored at - 80°C. Using a Bead Ruptor 12 (OMNI International), samples were homogenized with ∼6 ceramic beads (2.8 mm; OMNI International) at the highest speed for 2-3 rounds of 15 seconds. Chloroform extraction was performed on homogenized samples in TRIzol. RNA was isolated and purified using the RNeasy PLUS Mini Kit (Qiagen) according to manufacturer instructions. RNA concentration was quantified with a NanoDrop 2000 (ThermoFisher Scientific). 2500 ng of cDNA was synthesized per sample using SuperScript IV VILO Master Mix (ThermoFisher Scientific) and a C100 Touch Thermocycler (BioRad). 100 ng of cDNA was plated per well and the qRT-PCR reactions were performed on the StepOne Plus Real-Time PCR System (Applied Biosystems) using TaqMan Gene Expresion Master Mix (Applied Biosystems) according to manufacturer instructions. A list of TaqMan gene expression probes is detailed in SI Appendix, Table S2. Reference housekeeping genes include *Gapdh*, *B2m*, *Rer1*, and *Hprt*, as listed in the figure captions. Samples were normalized to the control (uninfected) group. All qRT-PCR data was analyzed using the 2^-ΔΔCt^ method (77).

### Tissue preparation for flow cytometry and analysis

#### Quadricep muscle with implant

Quadricep muscle tissues with implants (PCL, ECM, PEG, or HA hydrogel) were excised from the mouse, finely diced with a razor blade, and digested on a shaker for 45 minutes at 37°C with 1.67 Wünsch U/mL (5 mg/mL) of Liberase TL (Roche Diagnostics) and 0.2 mg/mL DNase I (Roche Diagnostics) in RPMI-1640 medium with L-Glutamine (Gibco) and 25 mM HEPES (Gibco). Digestion enzymes were neutralized by placing samples on ice and adding cold RPMI-1640 medium supplemented with L-Glutamine, 25 mM HEPES, and 1% bovine serum albumin (BSA) (Sigma). Tissues were mashed through 70 μm nylon strainers (Falcon) with excess 1X DPBS (Gibco), then filtered through 40 μm nylon strainers (Falcon) with excess 1X DPBS. Samples were spun down at 400xg for 5 minutes, resuspended with 1X DPBS, and transferred to a 96-well round bottom plate (Sarstedt). The entire cell pellet containing one quadricep muscle tissue with implant was plated per well.

#### Blood

Blood was collected from the submandibular vein of isoflurane-anesthetized mice into K_2_ EDTA collection tubes (BD). 200 μL of blood was transferred to a 96-well round bottom plate (Sarstedt) and spun down at 400xg for 5 minutes. Red blood cells were lysed with 100 μL of 1X PharmLyse^TM^ Lysing Buffer (BD) for 1 minute at room temperature. 150 μL of 1X DPBS (Gibco) was added to dilute the lysis buffer, and the samples were spun down at 400xg for 5 min. Samples were washed with 1X DPBS 4-5 more times until the liquid above the pellet was clear.

#### Spleen

Spleens were harvested from mice and mashed through a 70 μm nylon strainer (Falcon) with excess RPMI-1640 medium with L-Glutamine (Gibco) and 25 mM HEPES (Gibco). Samples were spun down at 400xg for 5 minutes, then lysed in 1 mL of 1X PharmLyse^TM^ Lysing Buffer (BD) for 1 minute at room temperature. Excess 1X DPBS (Gibco) was added and thoroughly mixed to dilute the lysis buffer, and the samples were spun down at 400xg for 5 minutes. The samples were then filtered through a 40 μm nylon strainer (Falcon), spun down at 400xg for 5 minutes, resuspended in 1X DPBS, and transferred to a 96-well round bottom plate (Sarstedt). Only ¼ to ½ of the spleen cell pellet was plated per well.

#### Bone marrow from femurs

Both femurs were harvested from mice and the surrounding muscle was scraped off. Femurs were placed in a 0.5 mL Eppendorf tube containing a hole at the bottom of the tube, which was nested in a 1.5 mL Eppendorf tube. Samples were spun at 10,000xg for 20 seconds and lysed in 1 mL of 1X PharmLyse^TM^ Lysing Buffer (BD) for 1 minute at room temperature. Excess RPMI-1640 medium with L-Glutamine (Gibco) and 25 mM HEPES (Gibco) was added and thoroughly mixed to dilute the lysis buffer, and samples were spun down at 300xg for 10 minutes. Samples were then filtered through a 70 μm nylon strainer (Miltenyi), spun down at 400xg for 5 minutes, resuspended in 1X DPBS, and transferred to a 96-well round bottom plate (Sarstedt). Only ½ of the cell pellet containing both femurs was plated per well (equivalent to 1 femur per well).

#### Inguinal lymph nodes

Both inguinal lymph nodes were harvested from mice and mashed through a 70 μm nylon strainer (Falcon) with excess RPMI-1640 medium with L-Glutamine (Gibco) and 25 mM HEPES (Gibco). Samples were spun down at 400xg for 5 minutes and transferred to a 96-well round bottom plate (Sarstedt). The entire cell pellet containing both inguinal lymph nodes was plated per well.

#### Surface marker staining

Once the samples were plated in the 96-well round bottom plate (Sarstedt), they were washed with 1X DPBS and stained with ZombieNIR Fixable Viability Stain (BioLegend) for 30 minutes on ice in the dark. Samples were washed twice with FACS buffer (1X DPBS [Gibco] supplemented with 1% BSA [Sigma] and 1 mM EDTA [Invitrogen]) and stained with surface marker antibodies that were prepared in FACS buffer supplemented with 1:20 TruStain FcX anti-mouse CD16/CD32 antibody (BioLegend), 1:50 Super Bright Complete Staining Buffer (eBioscience), and 1:50 True-Stain Monocyte Blocker (BioLegend) for 45 minutes on ice in the dark. Antibodies conjugated to certain protein-based tandem dyes (APC-Fire 750) were added to the antibody cocktail immediately before staining to reduce tandem dye aggregation. Viability and surface marker antibodies are listed in SI Appendix, Tables S3-S5. Samples were washed twice with FACS buffer, then fixed with 100 μL FluoroFix buffer (BioLegend) for 15 minutes at room temperature in the dark. Samples were washed twice with FACS buffer and resuspended in 1X DPBS. Prior to running the Aurora flow cytometer with the Automated Sample Loader (Cytek Biosciences), samples were spun down and resuspended in 1X DPBS. Data was unmixed with the SpectroFlo software (Cytek Biosciences) and gated according to the gating scheme in SI Appendix, Fig. S11.

#### Intracellular cytokine staining

Once the samples were plated in the 96-well round bottom plate (Sarstedt), cells were stimulated with 200 μL of 1X Cell Stimulation Cocktail (Plus Protein Transport Inhibitors) (eBioscience) in culture media supplemented with 5% fetal bovine serum (FBS) (Gibco) for 4 hours at 37°C. Samples were washed twice with 1X DPBS (Gibco) and stained with Fixable Viability Dye eFluor 780 (Invitrogen) for 30 minutes on ice in the dark. Samples were washed once with 1X DPBS and once with FACS buffer (1X DPBS [Gibco] with 5% FBS [Gibco]) supplemented with 1:50 TruStain FcX anti-mouse CD16/CD32 antibody (BioLegend) and 1:20 True-Stain Monocyte Blocker (BioLegend). Samples were stained with surface marker antibodies that were prepared in FACS buffer supplemented with 1:50 TruStain FcX anti-mouse CD16/CD32 antibody (BioLegend) and 1:20 True-Stain Monocyte Blocker (BioLegend) for 45 minutes on ice in the dark. Samples were washed twice with FACS buffer and fixed in 100 μL Cytofix/Cytoperm (BD) for 20 minutes on ice in the dark. Samples were washed twice with 1X Perm/Wash Buffer (BD) in distilled water and stained with intracellular marker antibodies that were prepared in 1X Perm/Wash Buffer for 30 minutes on ice in the dark. Samples were washed twice with 1X Perm/Wash buffer and resuspended in 1X DPBS before running on the Attune NxT flow cytometer (Thermo Fisher Scientific) the next day. Viability, surface marker, and intracellular marker antibodies are listed in SI Appendix, Table S6. Data was compensated on FlowJo v10.10.0 and gated according to the gating scheme in SI Appendix, Fig. S12.

#### Flow cytometry results

Flow cytometry results were presented as frequencies (percentage of larger parent population), calculated cell counts (total volume in well / volume acquired by cytometer * sample dilution factor * counts recorded by cytometer), or counts recorded by the cytometer using bar graphs depicting mean ± SD.

### Fluorescence activated cell sorting (FACS)

Tissue processing is the same as above with the following exceptions: 1) digestion enzymes were neutralized with cold RPMI-1640 medium supplemented with L-Glutamine, 25 mM HEPES, and 10% fetal bovine serum (FBS) (Gibco), 2) the viability stain used was Live/Dead Fixable Aqua (Invitrogen), and 3) there was no fixation step after surface marker staining. Viability and surface marker antibodies are listed in SI Appendix, Table S7. After surface staining, samples were washed twice with FACS buffer and resuspended in 1X DPBS (Gibco). Cell sorting was performed at the Johns Hopkins Sidney Kimmel Comprehensive Cancer Center High Parameter Flow Core using a BD FACSAria^TM^ Fusion Cell Sorter and BD FACSDiva software for data acquisition. Voltages and compensation were set using single stain cell controls and gates manually placed based on FMO controls. Fibroblasts (CD45^-^CD31^-^CD29^+^) from quadricep muscle tissue with PCL implant at 6wk post-implantation were sorted directly into 350 μL Buffer RLT Plus (Qiagen) with 1X beta-mercaptoethanol (Gibco). Samples were vortexed before freezing on dry ice and were stored at - 80°C.

### Bulk RNA sequencing

Sorted fibroblasts were submitted to the Johns Hopkins Single Cell & Transcriptomics Core for RNA extraction, QC, library prep, and bulk RNA sequencing. mRNA was isolated with the RNeasy Kit (Qiagen) and quantified with a NanoDrop spectrophotometer. QC was performed with the Agilent Fragment Analyzer. RNA samples had RQNs ranging from 9.3 to 10.0. cDNA synthesis and stranded mRNA Seq library preparation with unique dual indexes (UDI) was performed using Illumina Standard Total RNA Ligation Kit. QC was assessed with Qubit and Agilent Fragment Analyzer. Libraries were pooled and pair-end sequencing was performed using the Illumina NovaSeq platform with a SP 100 flow cell at a targeted depth of 80 million unique reads per sample. FASTQ files were aligned with STAR aligner (78) to the GENCODE release M27 (GRCm39) mouse genome assembly and annotation. Differential gene expression analysis was performed with DESeq2 (79) based on a negative binomial model with shrinkage estimation of the logarithmic fold change. Rank based gene set enrichment analysis for the REACTOME pathways gene sets were performed using the fgsea package (80), where differential gene expression results were ranked by logFC*-log10(p_adj_). Adjusted p values are reported based on the Benjamini-Hochberg method of multiple testing correction.

### Histological staining and imaging

#### Formalin fixation and paraffin embedding of tissues

Tissues were harvested and stored in 10% neutral buffered formalin (Sigma) for 48 hours. For the colon harvest, stool was removed via a syringe filled with 1X DPBS (Gibco) and tissues were Swiss-rolled before placing into 10% neutral buffered formalin (Sigma). Tissues were dehydrated through sequential steps in ethanols, cleared with xylene, and stored in paraffin overnight at 58-60°C. Tissues were embedded in paraffin blocks the following day. Embedded tissues were sectioned into 7 μm sections via microtome either in-house or through the Johns Hopkins Oncology Tissue Services SKCCC core facility.

#### Immunofluorescence

Citrullinated histone H3 (CitH3) (ab5103, Abcam) and Ly6G (ab238132, Abcam) were stained using the tyramide signal amplification (TSA) method with Opal 570 (Akoya Biosciences) and Opal 650 (Akoya Biosciences), respectively. Formalin-fixed paraffin-embedded (FFPE) sections of quadricep muscle tissue with PCL implant were baked for 1 hour at 58-60°C. Slides were deparaffinized and rehydrated through sequential steps in xylene, ethanols, and type I H_2_O. Slides were steamed for 15 minutes in antigen retrieval buffer (AR6, Akoya Biosciences), cooled to room temperature for 15-20 minutes, and rinsed with type I H_2_O. Peroxidases were quenched with 3% H_2_O_2_ (Sigma) in type I H_2_O for 15 minutes. Slides were blocked for 30 minutes with blocking buffer containing 10% BSA (Sigma) and 0.05% Tween 20 (Fisher Scientific) in 1X DPBS (Gibco), then incubated with 1:100 CitH3 primary antibody in blocking buffer for 30 minutes. Slides were washed 3 times in 1X Tris Buffered Saline with Tween 20 (TBST) (Cell Signaling Technology), followed by incubation with the secondary antibody, rabbit-on-rodent IgG HRP Polymer (Biocare Medical). Slides were washed 3 times in 1X TBST and incubated with 1:150 Opal 570 for 10 minutes. All steps from the Opal stain onwards were performed in the dark. Slides were washed 3 times in 1X TBST and the same steps as above (starting from the antigen retrieval step) were repeated for Ly6G, excluding the peroxidase quenching step. Ly6G was stained at a 1:500 concentration, and Opal 650 was used at 1:150. After the 3 washes in 1X TBST, slides were stained with spectral DAPI (Akoya Biosciences) for 5 minutes and rinsed 3 times with type I H_2_O. DAKO mounting medium (Agilent Technologies) was applied and slides were coverslipped (0.13-0.16 mm thickness, FisherScientific). Slides were dried overnight at room temperature in the dark and transferred to 4°C the next day. 3 days after staining, slides were imaged on a Zeiss Axio Imager A2 and stitched with the ZEN software.

#### Immunohistochemistry

FFPE sections of quadricep muscle tissue with PCL implant were baked for 1 hour at 58-60°C. Slides were deparaffinized and rehydrated through sequential steps in xylene, ethanols, and type I H_2_O. Slides were rinsed with 1X DPBS (Gibco) and tissues were circled with a PAP pen (Vector Laboratories). Slides were steamed in 1X citrate-based antigen unmasking solution (Vector Laboratories) for 25 minutes, then rinsed in 1X DPBS. Slides were incubated with hydrogen peroxide blocking agent for 20 minutes in the dark, washed twice with 1X DPBS, and incubated for 30 minutes in Protein Block (Abcam). Slides were washed twice with 1X DPBS and incubated overnight at 4°C with 1:100 aggrecan (AB1031, Chemicon) in DAKO Antibody Diluent. Slides were washed 3 times with 1X DPBS and incubated with biotinylated goat anti-rabbit IgG (Abcam) for 1 hour at room temperature. Slides were washed 3 times with 1X DPBS and AEC Substrate Kit (Abcam) was added for 4.5 minutes. (Incubation time was determined by monitoring the chromogenic change from aminoethyl carbazole [AEC] staining and using a primary delete control.) Slides were dipped in 1X DPBS, then type I H_2_O to stop incubation. Filtered Mayer’s hematoxylin (Abcam) was added to the slides for 10 seconds, and slides were rinsed in type I H_2_O until water ran clear. Slides were mounted with Aquamount (Lerner Laboratories) and coverslipped (0.13-0.16 mm thickness, FisherScientific). Slides were imaged with a digital slide scanner (Hamamatsu NanoZoomer-XR).

#### Picrosirius Red

FFPE sections of quadricep muscle with PCL implant were left in the oven for 15 minutes at 58- 60°C. Slides were stained with the Picro Sirius Red Stain Kit (Abcam) according to manufacturer protocol. DAKO mounting medium (Agilent Technologies) was applied and slides were coverslipped (0.13-0.16 mm thickness, FisherScientific). Slides were imaged with a polarizer on the Zeiss Axio Imager A2. Mean red-green values were measured via the analysis method depicted in SI Appendix, Fig. S7.

#### Masson’s Trichrome

FFPE sections of quadricep muscle tissue with PCL implant were stained by the Johns Hopkins Oncology Tissue Services SKCCC core facility. Slides were imaged with a digital slide scanner (Hamamatsu NanoZoomer-XR).

##### H&E

FFPE sections of Swiss-rolled colon tissue were stained by the Johns Hopkins Oncology Tissue Services SKCCC core facility. Slides were imaged with a digital slide scanner (Hamamatsu NanoZoomer-XR).

#### Atomic Force Microscopy (AFM) Fixed Microindentation

FFPE sections of quadricep muscle with PCL implant were deparaffinized and rehydrated through sequential steps in xylene, ethanols, and type I H_2_O. Slides were rehydrated in PBS overnight. Once placed on the AFM, slides were covered in PBS to facilitate measurements. Young’s Modulus was measured using a Nanowizard V BioScience (Bruker) equipped with SAA-SPH-1UM probes with a setpoint of 4 nN. Indentation tests were performed in the fibrotic tissue regions surrounding PCL particles. For each sample, 3-5 regions of size 50 x 50 µm were selected, with 64 measurements taken per selected region. The Young’s Modulus was calculated by fitting the force-displacement curves with the Hertz model with an assumed Poisson’s ratio of 0.5, with poorly fitting curves filtered out. For microrheology measurements, the probe oscillated in frequency shuffle of 1, 2.2, 4.5, 10, 22, 45, and 100 Hz for at least 5 cycles with an amplitude of 20 nm. The storage modulus, loss modulus, and loss factor were calculated by fitting the force-displacement oscillation curves with an assumed Poisson’s ratio of 0.5, with poorly fitting curves filtered out.

#### Statistics

All statistical analyses were performed with GraphPad Prism v10.1.2 (excluding bulk RNA sequencing analysis) with statistical significance defined as p≤0.05. Data were analyzed with a two-tailed unpaired t test assuming equal standard deviations or a 2-way ANOVA with Sidak’s multiple comparisons. The statistical analysis used for each graph is detailed in the figure captions.

## Acknowledgments

We would like to thank the Johns Hopkins Sidney Kimmel Comprehensive Cancer Center (SKCCC) High Parameter Flow Core for assistance with FACS. We thank Linda Orzolek, Jasmeet Sethi, and the Johns Hopkins Single Cell & Transcriptomics Core for assistance with RNA extraction, QC, library prep, and bulk RNA sequencing of sorted fibroblasts. We would like to thank Oncology Tissue Services Core SKCCC for sectioning FFPE tissues and staining them for Masson’s Trichrome and H&E. We would also like to thank Maria Browne for performing digital slide scanning of the Masson’s Trichrome and H&E-stained slides. We thank Integra for providing MicroMatrix UBM Particulate and funding the portion of the study related to this material. We thank Christopher K. Hee and Allergan for providing clinical hyaluronic acid hydrogels and funding the portion of the study related to these hydrogels. This study was funded by the NIH Pioneer Award DP1AR076959 (J.H.E.), Bloomberg∼Kimmel Institute for Cancer Immunotherapy (J.H.E., D.M.P., C.L.S.), Morton Goldberg Professorship (J.H.E.), Willowcroft Foundation (F.H.), Commonwealth Foundation (F.H.), Johns Hopkins University School of Medicine (C.L.S.), NIH Pathway to Independence Award K99AG081564 (J.C.M.), and the NSF Graduate Research Fellowship Program (B.Y. DGE2139757, A.R. DGE1746891, C.D.A. DGS2139754).

## Author Contributions

B.Y., N.R., A.R., E.G.G., F.H., D.M.P., C.L.S., J.H.E. designed research; B.Y., N.R., A.R., E.G.G., D.R.M., S.H.K., J.C.M., J.S.T.H., I.V., P.M., N.C., M.E., P.A., A.T. performed research; X.W., S.W., A.C., C.D.A., S.G., and A.T. contributed new reagents/analytical tools; B.Y., N.R., A.R., E.G.G., D.R.M., K.K., J.S.T.H., F.H., D.M.P., C.L.S., J.H.E. analyzed data; and B.Y., N.R., A.R., E.G.G., D.R.M., K.K., C.D.A., J.C.M., C.L.S., J.H.E. wrote the paper.

## Competing Interest Statement

J.H.E. holds equity in Unity Biotechnology and Aegeria Soft Tissue and is a consultant for Tessara. D.M.P. is a consultant at Aduro Biotech, Amgen, AstraZeneca, Bayer, Compugen, DNAtrix, Dynavax Technologies Corporation, Ervaxx, FLX Bio, Immunomic, Janssen, Merck, and Rock Springs Capital. D.M.P. holds equity in Aduro Biotech, DNAtrix, Ervaxx, Five Prime therapeutics, Immunomic, Potenza, Trieza Therapeutics. D.M.P. is a member of the scientific advisory board for Bristol Myers Squibb, Camden Nexus II, Five Prime Therapeutics, and WindMil. D.M.P. is a member of board of directors in Dracen Pharmaceuticals.

