## Supplementary Materials for "Murine gut microbiota dysbiosis via enteric infection modulates the foreign body response to a distal biomaterial implant"

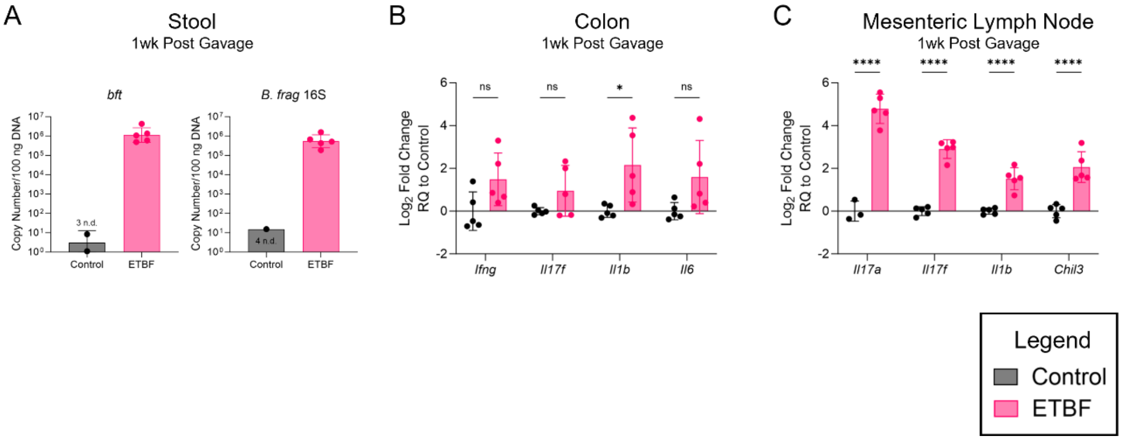


**Fig. S1**. Effect of ETBF infection on GI tissues. (A) Stool detection of *bft* and *B. fragilis 16S* at 1-week post gavage of DPBS or ETBF. Copy number calculated through standard curve. n.d. = not detected. (B) Colon expression of inflammatory genes at 1-week post ETBF, normalized to the control condition. *Gapdh* used as reference housekeeper gene. (C) Mesenteric lymph node expression of inflammatory genes at 1-week post ETBF, normalized to the control condition. *B2m* used as reference housekeeper gene. Two mice in control condition had Ct>35 for *Il17a* and were excluded from analysis. Graphs show geometric mean ± geometric SD (A) or mean ± SD (B-C). N=4-5 (A-C). *P < 0.05, **P < 0.01, ***P < 0.001, and ****P < 0.0001 by two-way ANOVA with Sidak's multiple comparisons (B-C).

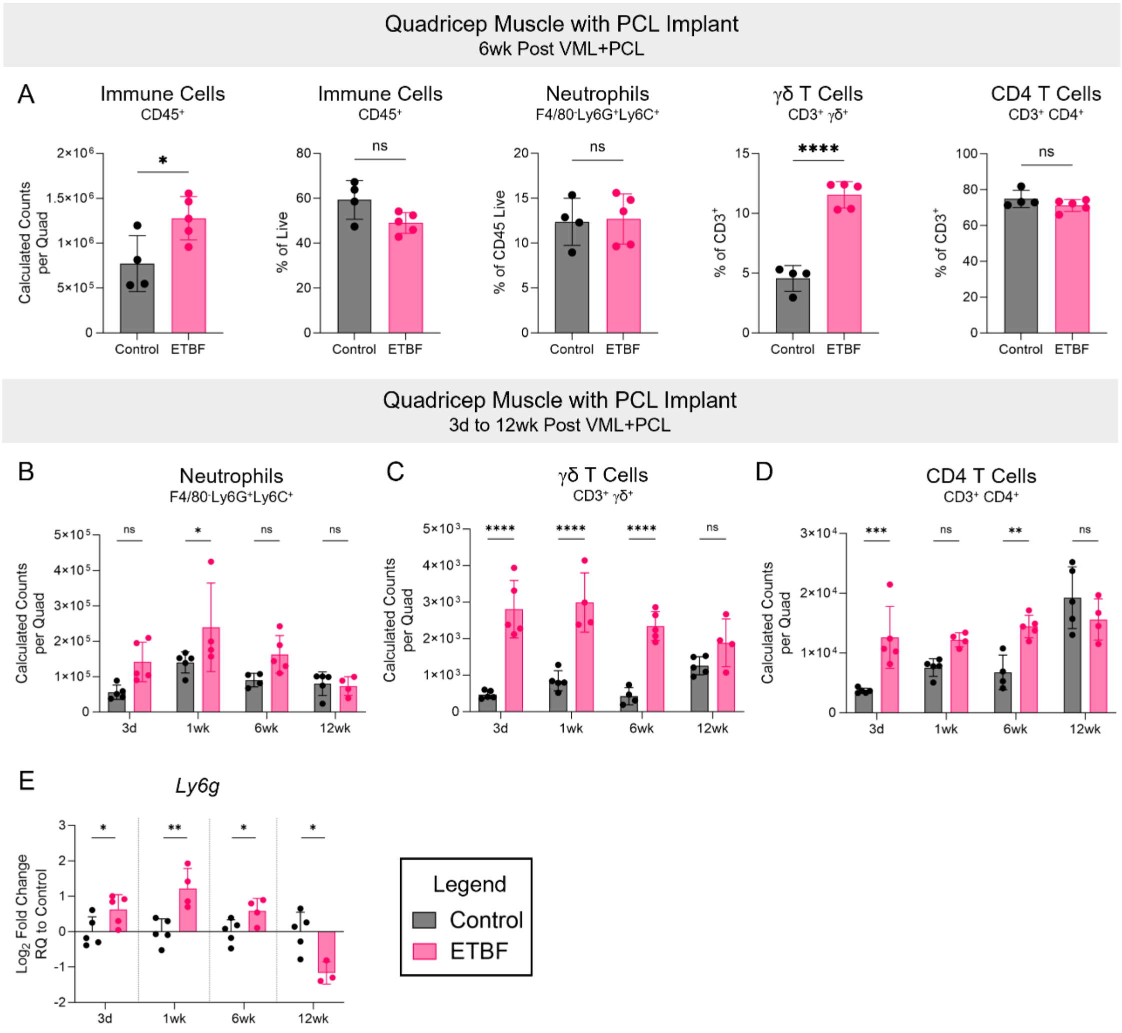


**Fig. S2.** Impact of ETBF infection on various immune cells at PCL implant site. (A) Calculated counts and/or frequency of immune cells ($CD45^+$), neutrophils ($CD11b^+F4/80^-Ly6G^+Ly6C^+$), γδ T cells ($CD3^+γδ^+$), and CD4 T cells ($CD3^+CD4^+$) in quadricep muscle with PCL implant at 6-weeks post VML+PCL in mice gavaged with DPBS or ETBF. Calculated counts of (B) neutrophils ($CD11b^+F4/80^-Ly6G^+Ly6C^+$), (C) γδ T cells ($CD3^+γδ^+$), and (D) CD4 T cells ($CD3^+CD4^+$) in quadricep muscle with PCL implant at 3-days, 1-week, 6-weeks, and 12-weeks post VML+PCL in mice gavaged with DPBS or ETBF. (E) *Ly6g* expression in PCL implant site at 3-days, 1-week, 6-weeks, and 12-weeks post VML+PCL in mice gavaged with DPBS or ETBF. Graphs show mean ± SD (A-E). N=3-5 (A-E). *$P < 0.05$, **$P < 0.01$, ***$P < 0.001$, and ****$P < 0.0001$ by two-way ANOVA with Sidak's multiple comparisons (B-D) or two-tailed unpaired t test (A, E).

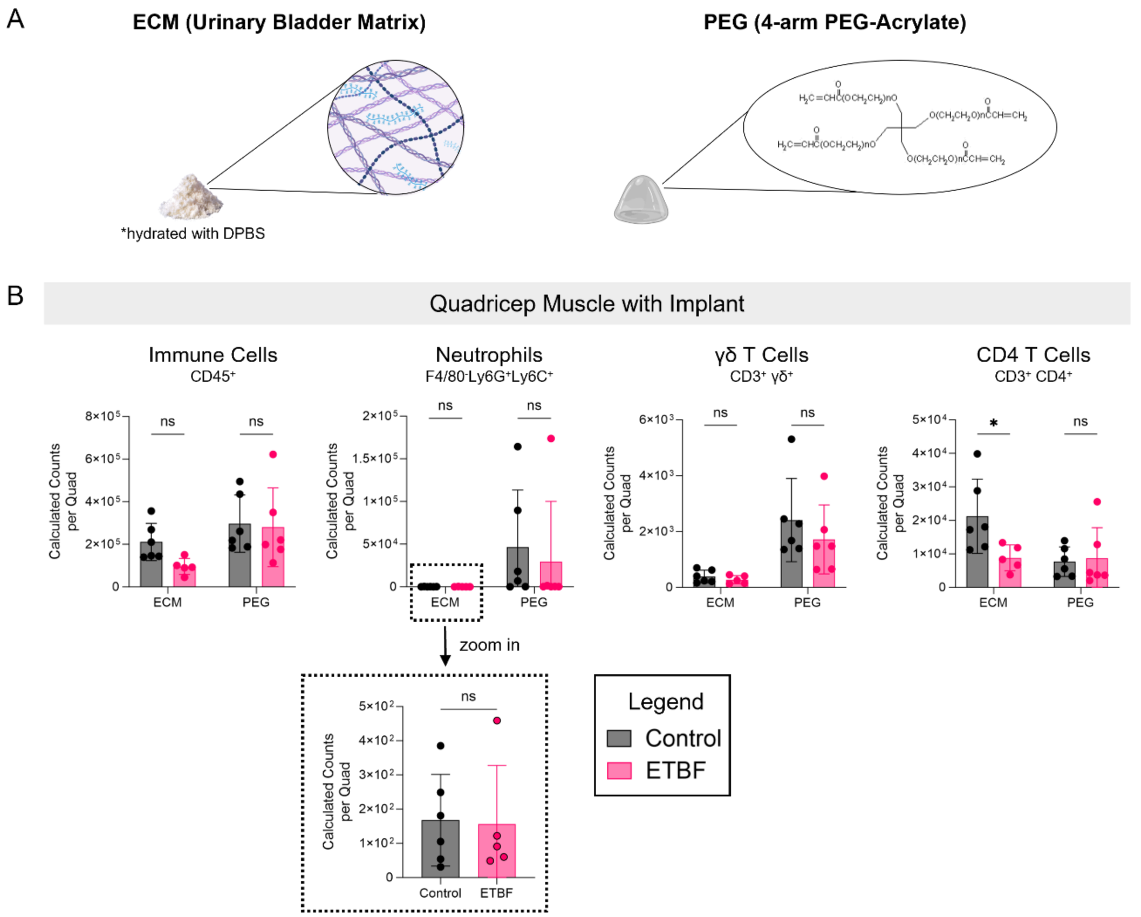


**Fig. S3**. Impact of ETBF infection on immune cell infiltration into ECM and PEG implant sites. (A) Description of ECM and PEG implants. ECM implant is porcine urinary bladder matrix from Integra and PEG implant is 4-arm PEG-acrylate hydrogel. Made with BioRender. (B) Calculated counts of immune cells ($CD45^+$), neutrophils ($CD11b^+F4/80^-Ly6G^+Ly6C^+$), γδ T cells ($CD3^+\gamma\delta^+$), and CD4 T cells ($CD3^+CD4^+$) in quadricep muscle with implant at 6-weeks post VML+implant in mice gavaged with DPBS or ETBF. Graphs show mean ± SD (B). N=5-6 (B). *P < 0.05, **P < 0.01, ***P < 0.001, and ****P < 0.0001 by two-way ANOVA with Sidak's multiple comparisons (B).

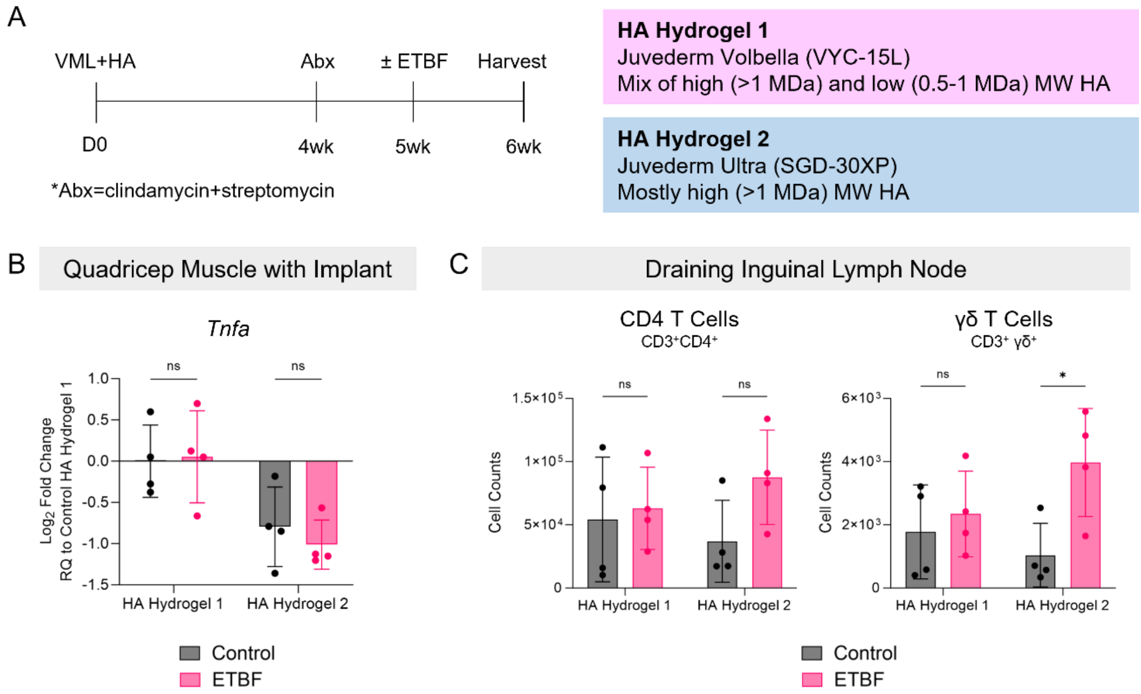


**Fig. S4**. Impact of ETBF infection on immune response to distal HA hydrogel implant. (A) Experimental timeline for data in Fig. S4 and description of the two hyaluronic acid hydrogels tested in this experiment. MW = molecular weight. (B) Quadricep muscle with HA hydrogel implant expression of *Tnfa* at 6-weeks post VML+PCL in mice gavaged with ETBF, normalized to the control HA hydrogel 1 condition. *Rer1* used as reference housekeeper gene. (C) Intracellular cytokine staining data of CD4 T cell ($CD3^{+}CD4^{+}$) and γδ T cell ($CD3^{+}γδ^{+}$) counts in draining inguinal lymph nodes of mice gavaged with DPBS or ETBF at 6-weeks post VML+HA hydrogel (n=4). Graphs show mean ± SD (B-C). N=4 (B-C). *P < 0.05, **P < 0.01, ***P < 0.001, and ****P < 0.0001 by two-way ANOVA with Sidak's multiple comparisons (B-C).

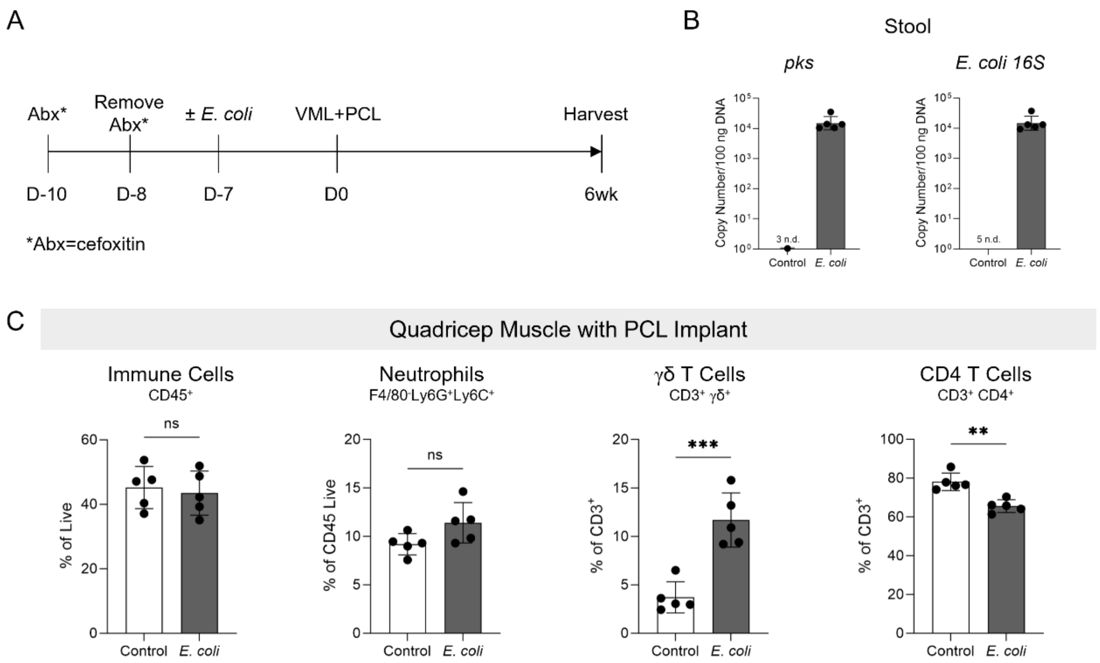


**Fig. S5**. Impact of *E. coli* infection on immune cell infiltration into distal PCL implant site. (A) Experimental timeline for data in Fig. S5. Cefoxitin was used instead of clindamycin and streptomycin for the antibiotic treatment, as the *pks*$^{+}$ *E. coli* strain used (see Methods) is resistant to cefoxitin and sensitive to streptomycin. (B) Stool detection of *pks* and *E. coli 16S* at the 6-week harvest time point. Copy number calculated through standard curve. n.d. = not detected. (C) Frequencies of immune cells (CD45$^{+}$), neutrophils (CD11b$^{+}$F4/80$^{-}$Ly6G$^{+}$Ly6C$^{+}$), γδ T cells (CD3$^{+}$γδ$^{+}$), and CD4 T cells (CD3$^{+}$CD4$^{+}$) in quadricep muscle with PCL implant at 6-weeks post VML+PCL in mice gavaged with DPBS or *E. coli*. Graphs show geometric mean ± geometric SD (B) or mean ± SD (C). N=5 (B-C). *P < 0.05, **P < 0.01, ***P < 0.001, and ****P < 0.0001 by two-tailed unpaired t test (C).

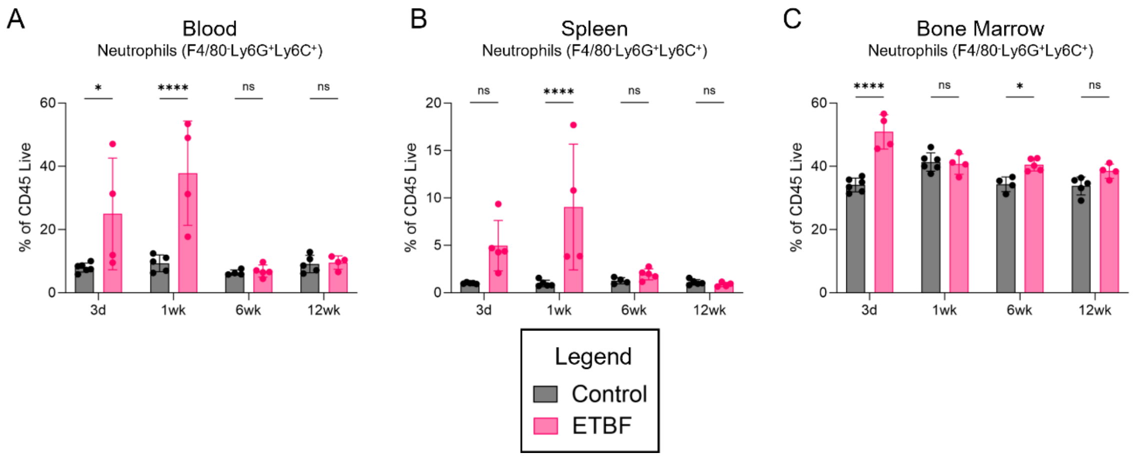


**Fig. S6**. Time course of impact of ETBF infection on neutrophil frequencies in systemic tissues. Neutrophil ($CD11b^+F4/80^-Ly6G^+Ly6C^+$) percentage of $CD45^+$ population in (A) blood, (B) spleen, and (C) bone marrow at 3-days, 1-week, 6-weeks, and 12-weeks post VML+PCL in mice gavaged with DPBS or ETBF. Graphs show mean ± SD (A-C). N=4-5 (A-C). *P < 0.05, **P < 0.01, ***P < 0.001, and ****P < 0.0001 by two-way ANOVA with Sidak's multiple comparisons (A-C).

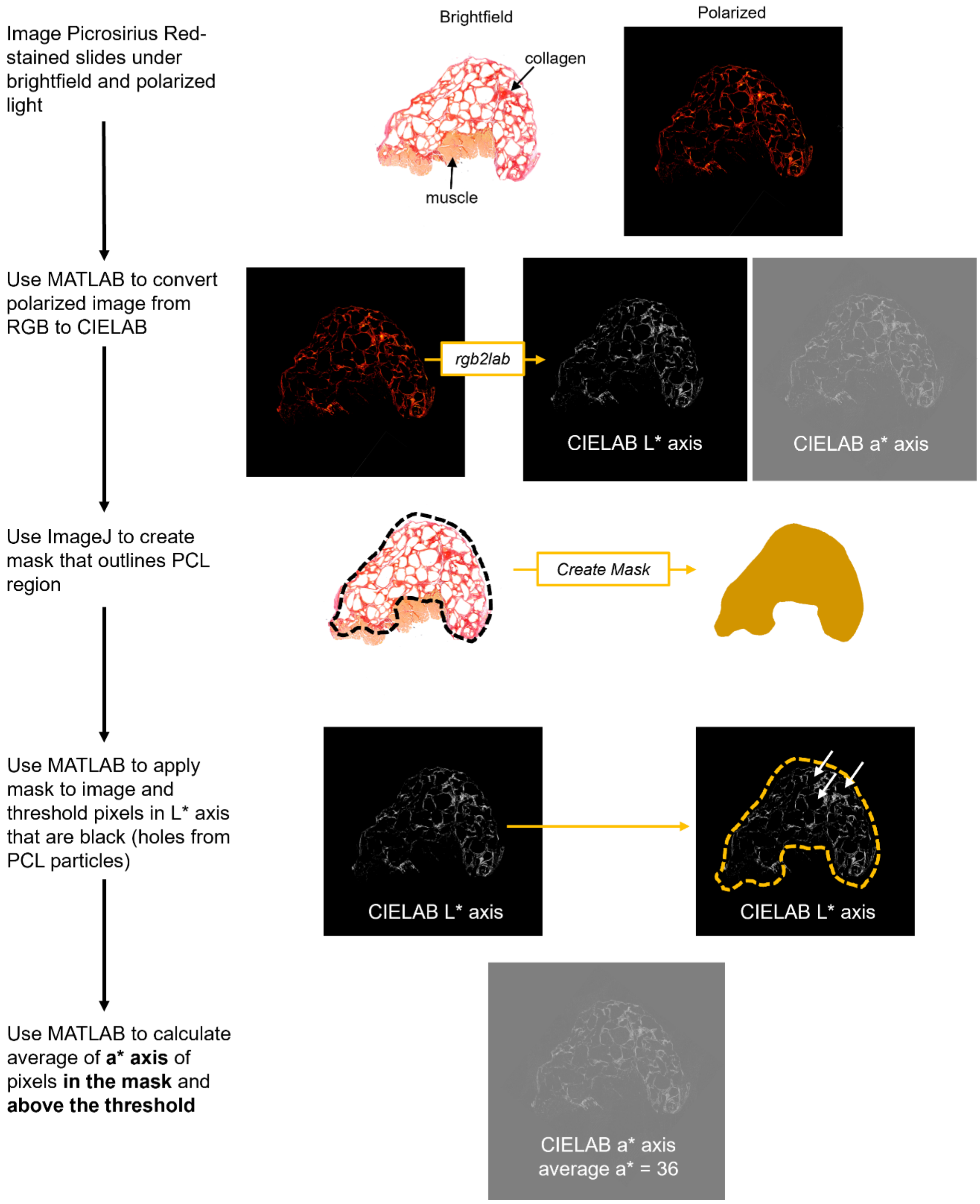


**Fig. S7**. Picrosirius Red analysis method.

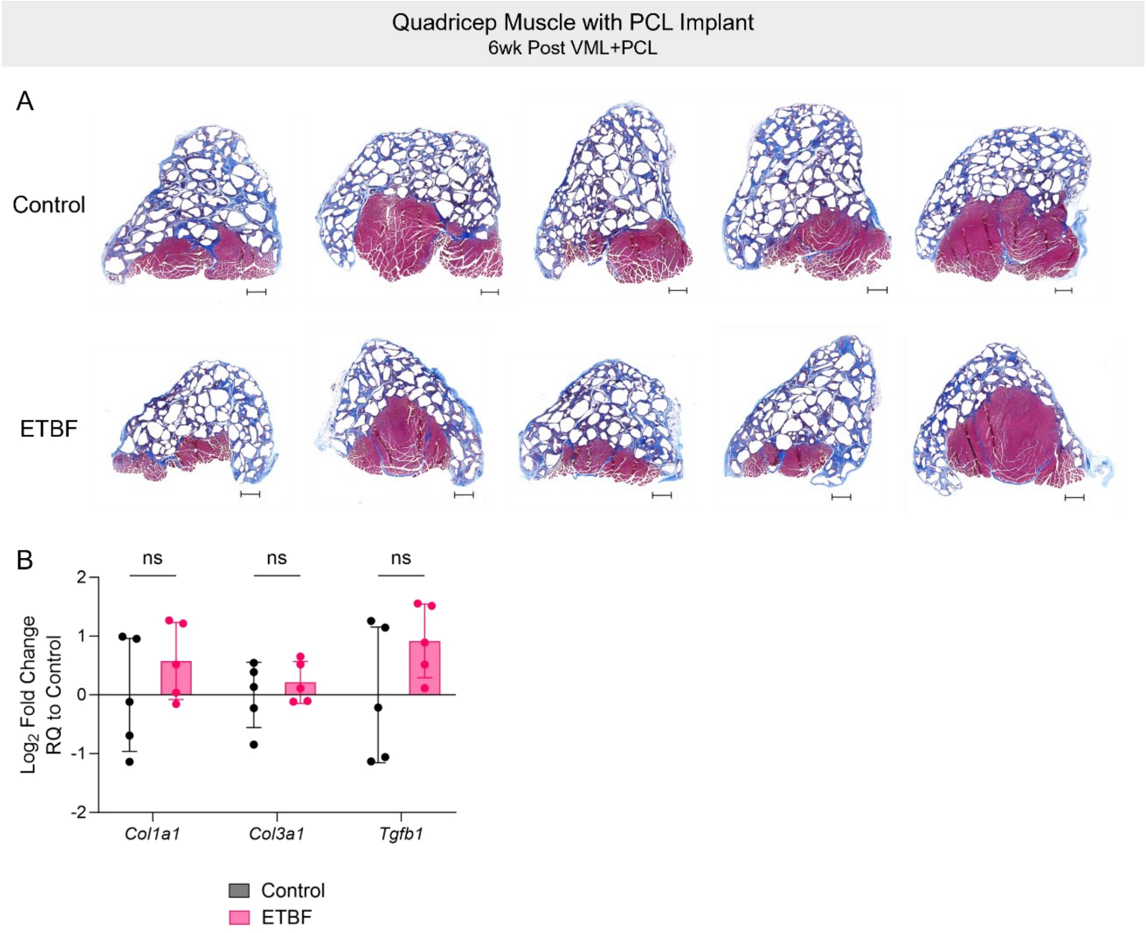


**Fig. S8**. Impact of ETBF infection on fibrosis readouts for PCL implant at 6-week time point. (A) Masson's trichrome images of quadricep muscle with PCL implant at 6 weeks post VML+PCL in mice gavaged with DPBS or ETBF. Scale bar, 500 µm. (B) Quadricep muscle with PCL implant expression of fibrosis-related genes at 6-weeks post VML+PCL in mice gavaged with ETBF, normalized to control condition. *Hprt* used as reference housekeeping gene. Graphs show mean ± SD. N=5 (B). *P < 0.05, **P < 0.01, ***P < 0.001, and ****P < 0.0001 by two-way ANOVA with Sidak's multiple comparisons (B).

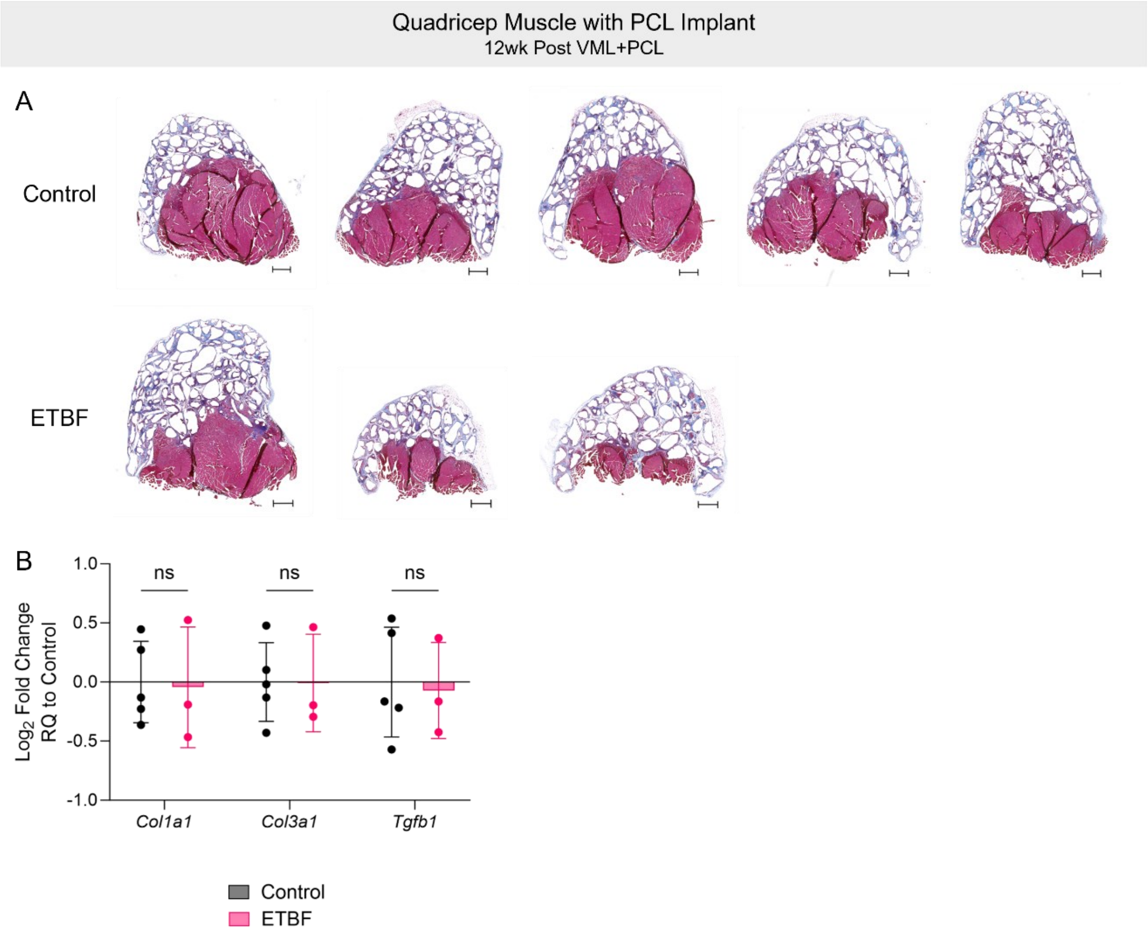


**Fig. S9**. Impact of ETBF infection on fibrosis readouts for PCL implant at 12-week time point. (A) Masson's trichrome images of quadricep muscle with PCL implant at 12-weeks post VML+PCL in mice gavaged with DPBS or ETBF. Scale bar, 500 µm. (B) Quadricep muscle with PCL implant expression of fibrosis-related genes at 12-weeks post VML+PCL in mice gavaged with ETBF, normalized to control condition. *Hprt* used as reference housekeeping gene. Graphs show mean ± SD. N=3-5 (B). *P < 0.05, **P < 0.01, ***P < 0.001, and ****P < 0.0001 by two-way ANOVA with Sidak's multiple comparisons (B).

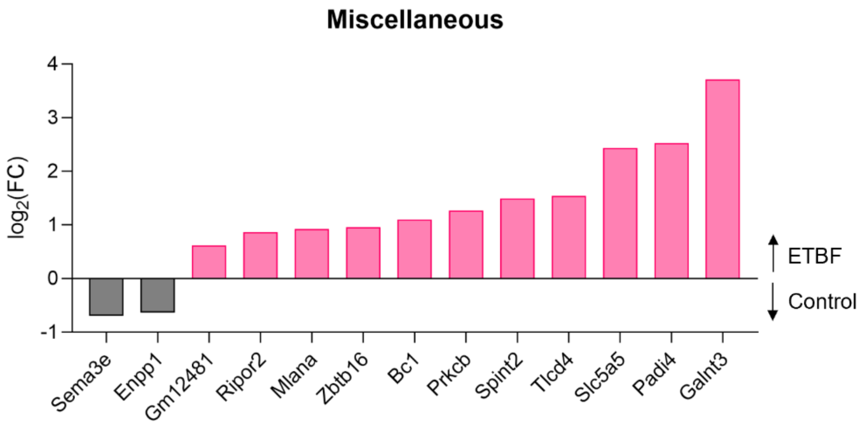


**Fig. S10**. Miscellaneous differentially expressed genes (DEGs) in sorted fibroblasts. Miscellaneous category of significant ($p_{adj} < 0.05$) DEGs in sorted fibroblasts ($CD45^-CD31^-CD29^+$) from quadricep muscle with PCL implant at 6-weeks post VML+PCL.

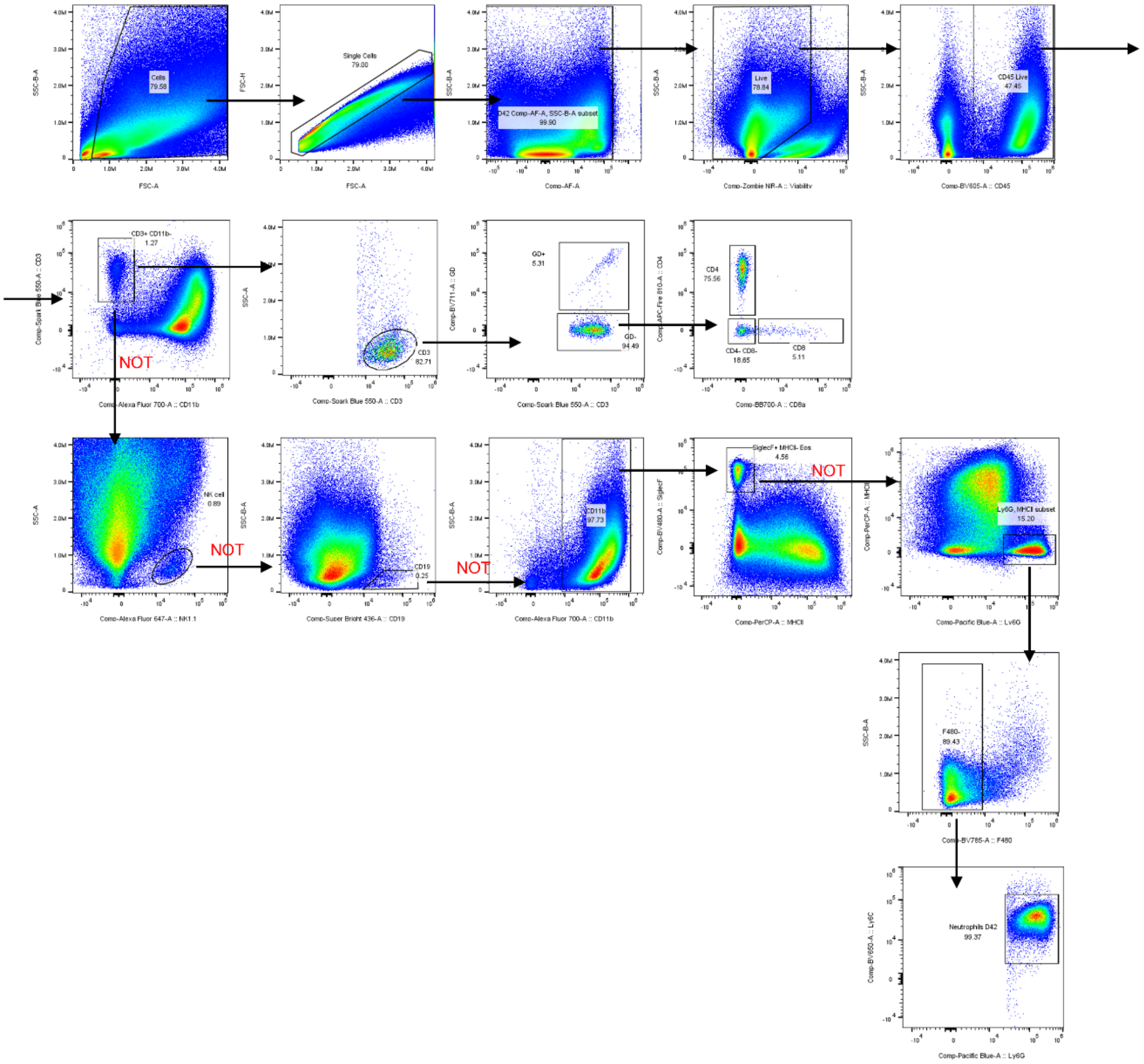

**Fig. S11**. Gating scheme for Cytek Aurora Pan Immune flow cytometry panel in quadricep muscle with PCL implant.

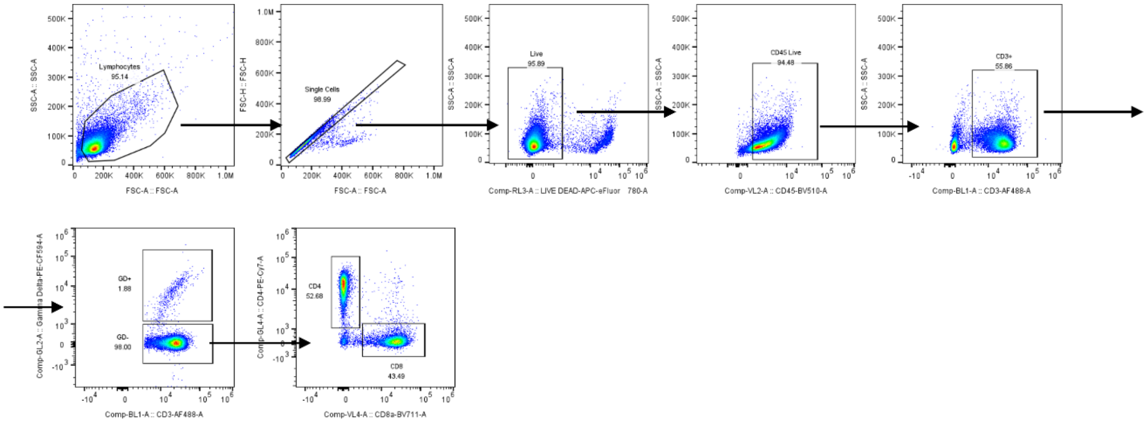


**Fig. S12**. Gating scheme for Attune NxT Intracellular Cytokine Staining flow cytometry panel in inguinal lymph nodes.

**Table S1.** Murine primers and probes for stool qPCR.

| Primer or Probe | Sequence (5' to 3') |
|---|---|
| *bft* F primer | GCGAACTCGGTTTATGCAGT |
| *bft* R primer | GTTGTAGACATCCCACTGGC |
| *bft* probe | 5'FAM - AGCAGAAGGTTATGACGA - 3' NFQ-MGB |
| *B. fragilis 16S* F primer | TCRGGAAGAAAGCTTGCT |
| *B. fragilis 16S* R primer | CATCCTTTACCGGAATCCT |
| *B. fragilis 16S* probe | 5'HEX - ACACGTATCCAACCTGCCCTTTACTCG - 3'BHQ1 |
| *pks* F primer | GCGCATCCTCAAGAGTAAATA |
| *pks* R primer | GCGCTCTATGCTCATCAACC |
| *pks* probe | 5'FAM - TATTCGACACAGAACAACGCCGGT - 3'BHQ1 |
| *E. coli 16S* F primer | CATGCCGCGTGTATGAAGAA |
| *E. coli 16S* R primer | CGGGTAACGTCAATGAGCAAA |
| *E. coli 16S* probe | 5'HEX - TCGGGTTGTAAAGTACTTTCAGCGGG - 3'BHQ1 |

**Table S2.** Murine TaqMan gene expression probes.

| Probe | Assay ID |
|---|---|
| *B2m* | Mm00437762_m1 |
| *Chil3* | Mm00657889_mH |
| *Col1a1* | Mm00801666_g1 |
| *Col3a1* | Mm01254477_m1 |
| *Gapdh* | Mm99999915_g1 |
| *Hprt* | Mm03024075_m1 |
| *Ifng* | Mm01168134_m1 |
| *Il17a* | Mm00439618_m1 |
| *Il17f* | Mm00521423_m1 |
| *Il1b* | Mm00434228_m1 |
| *Il6* | Mm00446190_m1 |
| *Ly6g* | Mm04934123_m1 |
| *Myf5* | Mm00435125_m1 |
| *Myod1* | Mm00440387_m1 |
| *Myog* | Mm00446194_m1 |
| *Nos2* | Mm00440502_m1 |
| *Pax7* | Mm01354484_m1 |
| *Rer1* | Mm00471276_m1 |
| *Tgfb1* | Mm01178820_m1 |
| *Tnfa* | Mm00443258_m1 |
| *Trem1* | Mm01278455_m1 |

**Table S3.** Cytek Aurora Pan Immune Flow Cytometry Panel for Quadricep Muscle with Implant

| Fluorophore | Antigen | Clone | Dilution | Manufacturer |
|---|---|---|---|---|
| BV421 | CD86 | GL-1 | 200 | BioLegend |
| SB436 | CD19 | 1D3 | 100 | ThermoFisher |
| Pac Blue | Ly6G | 1A8 | 250 | BioLegend |
| BV480 | SiglecF | E50-2440 | 100 | BD Biosciences |
| BV605 | CD45 | 30-F11 | 300 | BioLegend |
| BV650 | Ly6C | HK1.4 | 1200 | BioLegend |
| BV711 | γδ TCR | GL3 | 200 | BD Biosciences |
| BV785 | F4/80 | BM8 | 300 | BioLegend |
| SparkBlue550 | CD3 | 17A2 | 100 | BioLegend |
| PerCP | MHCII | M5/114.15.2 | 200 | BioLegend |
| BB700 | CD8a | 53-6.7 | 100 | BD Biosciences |
| PE | CD301b | URA-1 | 500 | BioLegend |
| PE-Dazzle594 | CD11c | N418 | 500 | BioLegend |
| APC | CD206 | C068C2 | 200 | BioLegend |
| AF647 | NK1.1 | PK136 | 200 | BioLegend |
| AF700 | CD11b | M1/70 | 400 | BioLegend |
| Zombie NIR | Viability | - | 5000 | BioLegend |
| APC-Fire750 | CD9* | MZ3 | 500 | BioLegend |
| APC-Fire810 | CD4 | GK1.5 | 100 | BioLegend |

*Added to antibody cocktail immediately before staining

**Table S4.** Cytek Aurora Pan Immune Flow Cytometry Panel for Spleen and Blood

| Fluorophore | Antigen | Clone | Dilution | Manufacturer |
|---|---|---|---|---|
| SB436 | CD19 | 1D3 | 100 | ThermoFisher |
| Pac Blue | Ly6G | 1A8 | 250 | BioLegend |
| BV480 | SiglecF | E50-2440 | 100 | BD Biosciences |
| BV605 | CD45 | 30-F11 | 300 | BioLegend |
| BV650 | Ly6C | HK1.4 | 1200 | BioLegend |
| BV711 | γδ TCR | GL3 | 200 | BD Biosciences |
| BV750 | B220 | RA3-6B2 | 200 | BioLegend |
| BV785 | F4/80 | BM8 | 300 | BioLegend |
| SparkBlue550 | CD3 | 17A2 | 100 | BioLegend |
| PerCP | MHCII | M5/114.15.2 | 200 | BioLegend |
| BB700 | CD8a | 53-6.7 | 100 | BD Biosciences |
| AF647 | NK1.1 | PK136 | 200 | BioLegend |
| AF700 | CD11b | M1/70 | 400 | BioLegend |
| Zombie NIR | Viability | - | 3000 (blood)<br>5000 (spleen) | BioLegend |
| APC-Fire810 | CD4 | GK1.5 | 100 | BioLegend |

**Table S5.** Cytek Aurora Pan Immune Flow Cytometry Panel for Bone Marrow

| Fluorophore | Antigen | Clone | Dilution | Manufacturer |
|---|---|---|---|---|
| SB436 | CD19 | 1D3 | 100 | ThermoFisher |
| Pac Blue | Ly6G | 1A8 | 250 | BioLegend |
| BV605 | CD45 | 30-F11 | 300 | BioLegend |
| BV650 | Ly6C | HK1.4 | 1200 | BioLegend |
| BV750 | B220 | RA3-6B2 | 200 | BioLegend |
| BV785 | F4/80 | BM8 | 300 | BioLegend |
| PerCP | MHCII | M5/114.15.2 | 200 | BioLegend |
| AF700 | CD11b | M1/70 | 400 | BioLegend |
| Zombie NIR | Viability | - | 3000 | BioLegend |

**Table S6.** Attune NxT Intracellular Cytokine Staining Panel for Inguinal Lymph Nodes

| **Fluorophore** | **Antigen** | **Clone** | **Dilution** | **Manufacturer** |
|---|---|---|---|---|
| BV421 | FoxP3 | MF-14 | 150 | BioLegend |
| BV510 | CD45 | 30-F11 | 100 | BD |
| BV605 | NK1.1 | PK136 | 200 | BioLegend |
| BV711 | CD8a | 53-6.7 | 200 | BioLegend |
| AF488 | CD3 | GK1.5 | 150 | BioLegend |
| PerCP-Cy5.5 | CD19 | 6D5 | 250 | BioLegend |
| PE | IL-4 | 11B11 | 150 | BioLegend |
| PE/Dazzle 594 | γδ TCR | GL3 | 200 | BioLegend |
| PE-Cy7 | CD4 | GK1.5 | 250 | BioLegend |
| APC | IFNγ | XMG1.2 | 150 | BioLegend |
| AF700 | IL-17a | TC11-18H10.1 | 250 | BioLegend |
| eFluor780 | Viability | - | 1000 | ThermoFisher |

**Table S7.** FACS Panel for Bulk RNA Sequencing

| Fluorophore | Antigen | Clone | Dilution | Manufacturer |
|---|---|---|---|---|
| BV421 | CD31 | 390 | 200 | BioLegend |
| BV510 | Viability Aqua | - | 500 | Invitrogen |
| BV605 | CD45 | 30-F11 | 150 | BioLegend |
| BV711 | CD11b | M1/70 | 600 | BioLegend |
| AF488 | CD3 | 17A2 | 50 | BioLegend |
| PE-Cy7 | F4/80 | BM8 | 250 | BioLegend |
| APC-Cy7 | CD29 | HMB1-1 | 400 | BioLegend |
